# A systematic CRISPR screen reveals an IL-20/IL20RA-mediated peritoneal immune crosstalk to prevent the ovarian cancer metastasis

**DOI:** 10.1101/2021.01.03.425154

**Authors:** Jia Li, Xuan Qin, Jie Shi, Xiaoshuang Wang, Tong Li, Mengyao Xu, Xiaosu Chen, Yujia Zhao, Jiahao Han, Yongjun Piao, Wenwen Zhang, Pengpeng Qu, Longlong Wang, Rong Xiang, Yi Shi

## Abstract

Transcoelomic spread of cancer cells across the peritoneal cavity occurs in most initially diagnosed ovarian cancer (OC) patients and accounts for most cancer-related death. However, how OC cells interact with peritoneal stromal cells to evade the immune surveillance remains largely unexplored. Here, through an *in vivo* genome-wide CRISPR/Cas9 screen, we identified IL20RA, which decreased dramatically in OC patients during peritoneal metastasis, as a key factor preventing the transcoelomic metastasis of OC. Reconstitution of IL20RA in highly metastatic OC cells greatly suppresses the transcoelomic metastasis. OC cells, when disseminate into the peritoneal cavity, greatly induce peritoneum mesothelial cells to express IL-20 and IL-24, which in turn activate the IL20RA downstream signaling in OC cells to produce mature IL-18, eventually resulting in the polarization of macrophages into the M1-like subtype to clear the cancer cells. Thus, we show an IL-20/IL20RA-mediated crosstalk between OC and mesothelial cells that supports a metastasis-repressing immune microenvironment.

## Introduction

Ovarian cancer (OC) has the highest mortality among all gynecological malignancies that seriously threatens women’s health worldwide (Siegel et al., 2020). Among OC, epithelial ovarian cancer (EOC) is the most common type accounting for 90% of all cases(Farley et al., 2008). EOC is classified by tumor cell histology as serous (52%), endometrioid (10%), mucinous (6%), clear cell (6%), and other rare subtypes. Due to the lack of clear symptoms at the early stage, the spread of cancer cells across the peritoneal cavity occurs in most patients at the initial diagnosis, with approximately 70% of OC patients already at metastatic stage (stage III and stage IV)(Vaughan et al., 2011). For OC patients with metastasized cancer cells, current therapies including chemotherapy and targeted therapies can only achieve very limited clinical outcome, with the 5-year survival rate of 20-40% for OC patients at advanced stages(Torre et al., 2018).

Although OC cells can metastasize to other distant organs through blood vessels and lymphatic vessels, the transcoelomic metastasis of OC occurs most commonly, which includes multiple processes, such as detachment of tumor cells from primary sites, immune evasion of disseminated tumor cells in the peritoneal cavity, colonization of tumor cells on the omentum and peritoneum(Tan et al., 2006). Disseminated OC cells have to survive in a peritoneal environment constituted by lymphocytes, macrophages, natural killer (NK) cells, fibroblasts, methothelial cells, as well as cytokines and chemokines secreted by these cells for a successful transcoelomic metastasis(Ahmed & Stenvers, 2013; Worzfeld et al., 2017). Although the role of the immune microenvironment in peritoneal cavity is indisputable in the metastasis of OC, the molecular network regulating the crosstalk between the disseminated tumors cells and immune microenvironment in peritoneal cavity is still just a tip of the iceberg. Therefore, it is imperative to understand the specific immune microenvironment in the peritoneal cavity for developing more efficient immunotherapeutic strategies against metastatic OC.

To systematically identify key genes involved in the regulation of the peritoneal metastasis of OC, we performed genome-wide gene knockout screening in an orthotopic mouse model of OC by using CRISPR/Cas9 knockout library, which has been successfully utilized to discover novel genes in the occurrence and development of diseases, especially in cancers(Huang et al., 2019; Ng et al., 2020; Shalem et al., 2014). We identified interleukin 20 receptor subunit alpha (IL20RA) as a potent suppressor of the transcoelomic metastasis of OC. IL20RA is mainly expressed in epithelial cells and forms the functional receptor, when heterodimerized with interleukin 20 receptor subunit beta (IL20RB), to bind immune cell-produced IL-20 subfamily of cytokines IL-19, IL-20 and IL-24, that have essential roles in regulating epithelial innate immunity and tissue repair(Rutz et al., 2014). IL20RA and IL20RB are also detected in tumors of epithelial origin including breast cancer, non-small-cell lung cancer, and bladder cancer, while IL-20 subfamily of cytokines have been reported to have either tumor-promoting or tumor-suppressing roles depending on tumor types and the local immune environment(Gopalan et al., 2007; Lee et al., 2013; Pestka et al., 2004; Rutz et al., 2014; Whitaker et al., 2012). In the present study, we discovered a novel IL-20/IL20RA-mediated crosstalk between disseminated OC cells and peritoneum mesothelial cells that eventually promotes the generation of M1-like inflammatory macrophages to prevent peritoneal dissemination of OC cells.

## Results

### *IL20RA* is an anti-transcoelomic metastasis gene in OC identified by a genome-scale knockout screening *in vivo*

To screen key genes regulating transcoelomic metastasis of OC, we utilized human epithelial OC cell SK-OV-3 with relatively low metastatic capacity to set up the orthotopic transplant tumor model in NOD-SCID mice. Prior to the inoculation of these cells into the mouse ovaries, SK-OV-3 cells were transduced with the human CRISPR knockout library (GeCKO v2.0) made by Feng Zhang *et al*, which contains sgRNAs specifically against 19,050 protein-encoding genes and 1,864 miRNA genes and 1,000 non-targeting control sgRNAs(Shalem et al., 2014). Peritoneal metastasized SK-OV-3 cells in the intraperitoneal cavity were isolated and expanded *in vitro* for next runs of orthotopic transplant (**Fig. 1A**). After 3 runs of *in vivo* screening, highly metastatic SK-OV-3 cells were subjected to high-throughput sgRNA library sequencing to reveal the sgRNA representations. The RNAi Gene Enrichment Ranking (RIGER) P value analysis was used to identify significantly enriched sgRNAs in metastasized (sgRNA^Met^) or primary (sgRNA^Pri^) OC xenografts. By using the criteria of the number of enriched sgRNAs targeting each gene ≥ 3, P-value < 0.05 and the normalized enrichment score (NES) < −1.2, we got two high-ranked genes, namely *IL20RA* and *TEX14* (**Fig. 1B**). Given all the 6 sgRNAs targeting IL20RA in GeCKO library were enriched with high NES, we chose IL20RA for further investigation on its function in the transcoelomic metastasis of OC.

**Figure 1.**
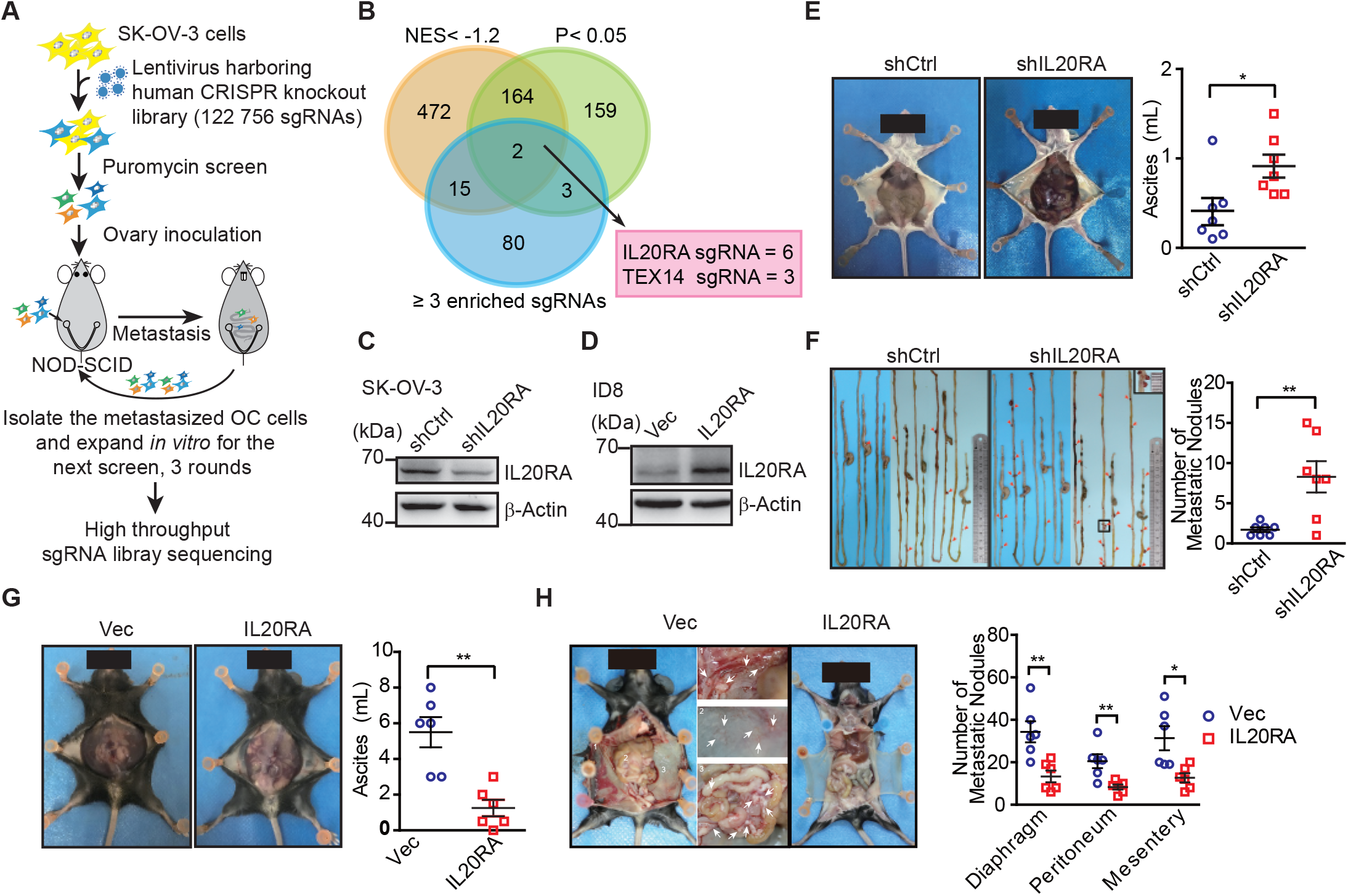
High-Throughput CRISPR screen identified IL20RA as a suppressor of the transcoelomic metastasis of OC. **A**. Schematics of experiment design to screen metastasis-related genes using CRISPR/Cas9 library in OC orthotopic murine model. **B**. Venn diagram comparing the hits met the indicated enrichment criteria. **C-D**. Western blot analysis of IL20RA in control shRNA (shCtrl)- or IL20RA shRNA (shIL20RA)-transfected SK-OV-3 cells (**C**) and in IL20RA- or empty vector-transfected ID8 cells (**D**). **E-F**. Representative images of NOD-SCID mice with shCtrl- or shIL20RA-transfected SK-OV-3 cells orthotopically transplanted in the ovaries at 40 days post-inoculation (E, left panel). The ascites volumes (**E**, right panel) and the numbers of metastatic nodules on the surfaces of intestines (**F**) were quantified (n = 7, data are shown as means ± SEM, *P<0.05, **P < 0.01, by unpaired two-sided student’s t-test). **G-H**. Representative images of C57BL/6 mice at 40 days after orthotopically inoculated with IL20RA-reconstituted or control ID8 cells in ovaries (**G**, left panel). The ascites volumes (**G**, right panel) and the numbers of metastatic nodules on the surfaces lining the peritoneal cavities (**H**) were quantified (n = 6, data are shown as means ± SEM, *P<0.05, **P < 0.01 by unpaired two-sided student’s t-test).

To confirm the role of IL20RA in OC metastasis, we knocked down IL20RA in SK-OV-3 cells, which had relatively high level of IL20RA (**Fig. 1C** and **Fig. S1, A and B**), and reconstituted IL20RA in highly metastatic murine epithelial OC cell ID8, which was isolated from peritoneal ascites of ID8-injected C57BL/6 mice as described by Kristy K Ward and colleagues(Ward et al., 2013) with high-grade serous ovarian carcinoma (HGSOC)-like characteristics(Diaz Osterman et al., 2019) and had very low level of endogenous IL20RA (**Fig. S1A** and **Fig. 1D**). In the SK-OV-3 orthotopic mouse OC model, silencing IL20RA greatly promoted the transcoelomic metastasis of OC and the formation of ascites (**Fig. 1, E and F**), while in ID8 syngeneic mouse OC model, reconstitution of IL20RA in ID8 cells dramatically reduced the volumes of ascites and the numbers of metastatic nodules in diaphragm, peritoneum and mesentery (**Fig. 1, G and H**).

To further investigate whether IL20RA suppresses transcoelomic metastasis of OC at the later stage when OC cells have disseminated into the peritoneal cavity, we directly injected ID8 cells into the peritoneal cavity of C57BL/6 mice. We were still able to observe that IL20RA-reconstituted ID8 cells formed much less metastases on the diaphragm, peritoneum and mesentery and resulted in much less ascites as well (**Fig. S1, C and D**). Together, these data suggest that IL20RA is a potent suppressor gene for the transcoelomic metastasis of OC.

### The IL20RA expression is dramatically decreased in metastases of OC patients and positively correlates with the clinical outcome

To further get the clinical evidences on the relevance of IL20RA in OC metastasis, we collected primary OC tissues and paired cancer cells isolated from ascites and metastatic nodules in peritoneal cavity from twenty serous OC patients. Analysis of IL20RA protein by western blot shows a dramatic decrease of IL20RA in the transcoelomic spread cancer samples when compared with those from the primary sites (**Fig. 2, A and B**), which is confirmed by the quantitative reverse-transcriptase-PCR (qRT-PCR) analysis of *IL20RA* mRNA (**Fig. 2, C and D**). Immunohistochemical (IHC) analysis also shows the much lower level of IL20RA in metastatic cancer cells than that in the cancer cells at the primary sites (**Fig. 2, E and F**). To further investigate the relevance of IL20RA with OC metastasis, we analyzed the correlation between the IL20RA level and the survival of serous OC patients given that metastasis accounts for over 90% of cancer death. High level of IL20RA significantly correlates with the better overall survival (OS) and progression-free survival (PFS) of serous OC patients (**Fig. 2G**), supporting that IL20RA functions as a suppressor of OC metastasis.

**Figure 2.**
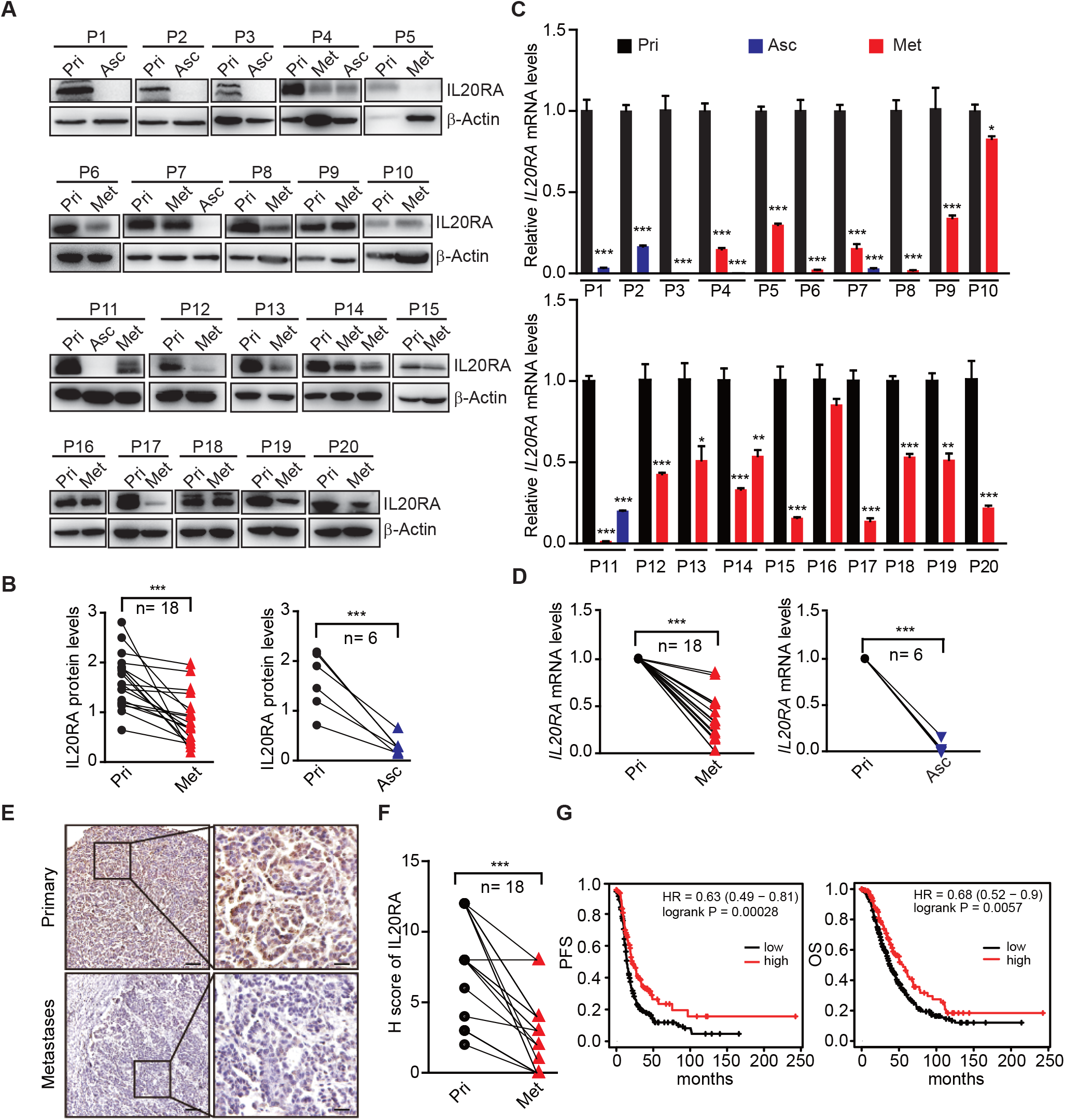
Dramatically decreased IL20RA in human OC peritoneal metastases and its correlation with the clinical outcome. **A-B.** Western blot analysis of IL20RA in primary OC tissues (Pri) and paired metastatic cancer cells in ascites (Asc) and metastatic nodules on the surfaces of the abdominal organs (Met) (**A**). Quantification results are plotted in (**B**) (n = 20, ***P < 0.001, by unpaired two-sided student’s t-test). **C-D**. qRT-PCR analysis of IL20RA in human primary OC tissue (Pri) and paired peritoneal metastases (Asc and Met) (**C**, data are plotted as means ± SEM from three independent measurements, *P < 0.05, **P < 0.01, ***P < 0.001, by unpaired two-sided student’s t-test). The comparison of the IL20RA levels in these two groups are analyzed in (**D**, **P < 0.01, ***P < 0.001, by unpaired two-sided student’s t-test). **E-F**. Representative images of IHC analysis of IL20RA in in human primary OC tissue and paired peritoneal metastases (**E**) and quantification by H-score (**F**, n = 18, ***P < 0.001, by paired two-sided student’s t-test). Scale bar: 100 μm (left panel in E); 20 μm (right panel in E). **G**. Kaplan-Meier survival plot to show the progression free survival (PFS) and overall survival (OS) of serous OC patients with different IL20RA expression.

By contrast, as a heterodimerization partner for IL20RA, IL20RB expression shows no significant difference between primary and metastatic OC specimen (**Fig. S2, A and B**) and negatively correlates with the OS and PFS of OC patients (**Fig. S2C**), suggesting that IL20RA is the key subunit of IL20RA/IL20RB receptor complex that is regulated during the transcoelomic metastasis of OC.

### Characterization of IL20RA-mediated modulation on the peritoneal immune microenvironment during the transcoelomic metastasis of OC

To get insights into the mechanisms of IL20RA-mediated inhibition on OC metastasis, we firstly excluded the possibility that IL20RA might regulate the proliferation and migration of OC cells (**Fig. S3A-C**). We next examined the possible impacts of IL20RA in shaping the immune microenvironment during the spreading of OC cells into the intraperitoneal cavity. In the syngeneic murine OC model by orthotopic transplant of ID8 cells, we analyzed the proportions of immune cells in ascites, including macrophages, T lymphocytes, and B lymphocytes. Compared with control ID8 cells with very low level of endogenous IL20RA (**Fig. S1A**), reconstitution of IL20RA does not change the total proportion of macrophages (CD11b^+^ F4/80^+^) among leukocytes (CD45^+^).

However, we observed a significant increase in the proportion of M1-like (MHC Ⅱ ^+^ CD206^−^) macrophages and a dramatic decrease of M2-like (MHC Ⅱ ^−^ CD206^+^) macrophages in the malignant ascites caused by IL20RA-reconstituted ID8 cells (**Fig. 3, A and B**), which were further confirmed by dramatically increased M1-like marker genes and decreased M2-like markers (**Fig. 3C**).

**Figure 3.**
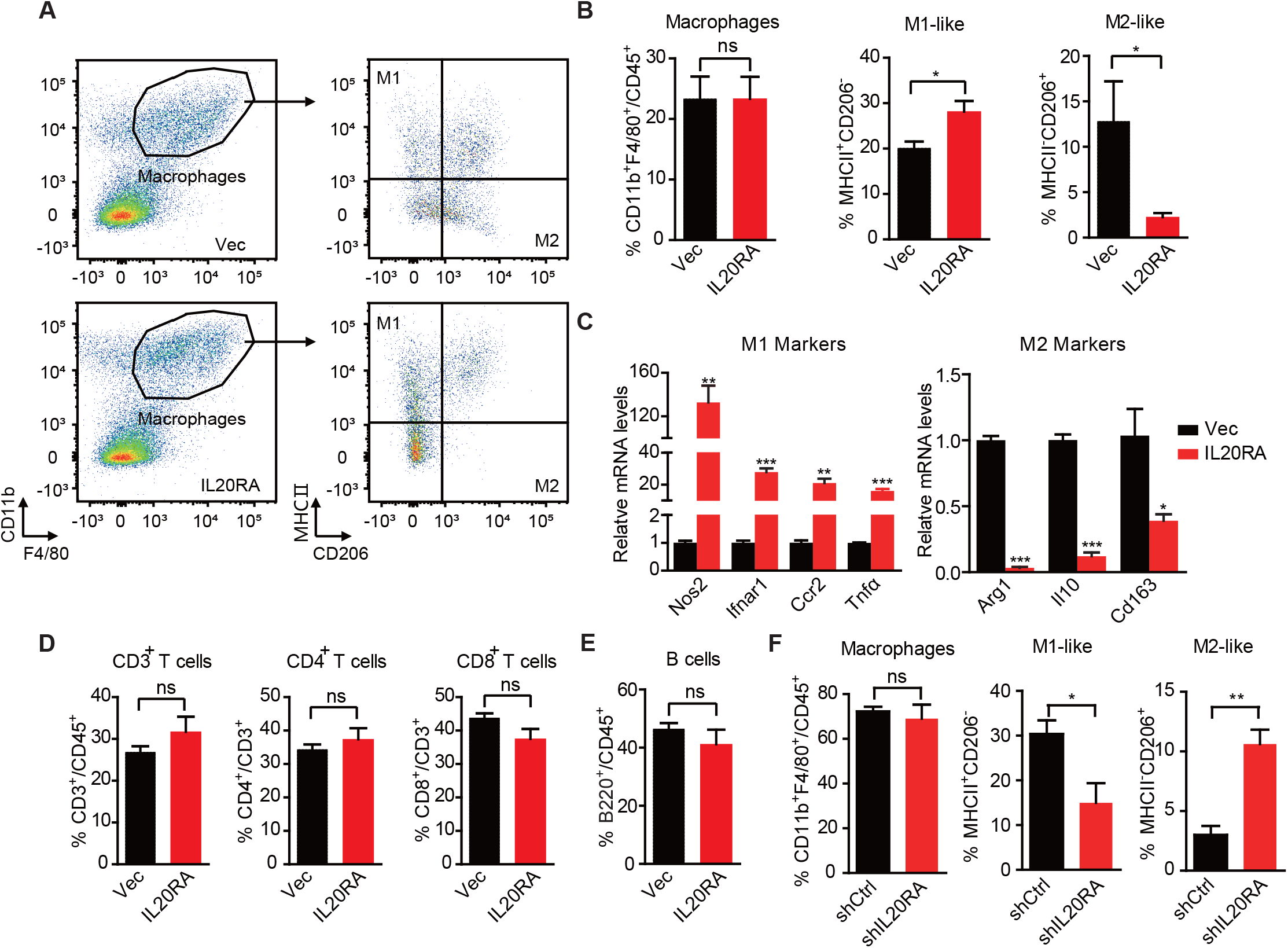
Characterization of IL20RA-mediated modulation on the peritoneal immune microenvironment during the transcoelomic metastasis of OC. **A-B.** Flow cytometry analysis of macrophages (CD45^+^ CD11b^+^ F4/80^+^) and M1-like (MHCⅡ^+^ CD206^−^) and M2-like (MHCⅡ^−^ CD206^+^) subpopulations in ascites formed in C57BL/6 mice at 60 days after orthotopically inoculated with IL20RA-reconsitituted or control (Vec) ID8 cells (**A**). The quantification is shown in (**B**) as means ± SEM (n = 5), *P < 0.05, ns not significant, by unpaired two-sided student’s t-test. **C**. qRT-PCR analysis of key marker genes in peritoneal macrophages (CD11b^+^ F4/80^+^) isolated in (**A**) (shown as means ± SEM, *P < 0.05, **P < 0.01, ***P < 0.001, by unpaired two-sided student’s t-test). **D-E**. Flow cytometry analysis of peritoneal T lymphocytes (**D**) and B lymphocytes (**E**) from syngeneic OC mouse model (same mice as described in **A**), ns not significant, by unpaired two-sided student’s t-test. **F**. Flow cytometry analysis of macrophages in ascites formed in NOD-SCID mice at 40 days after orthotopically inoculated with indicated SK-OV-3 cells. Data are shown as means ± SEM, n = 3, *P < 0.05, **P < 0.01, ***P < 0.001, by unpaired two-sided student’s t-test.

We also found that neither the ratios of T lymphocytes (CD45^+^ CD3^+^) and their subtypes (i.e., CD4^+^ and CD8^+^ T lymphocytes) nor that of B lymphocytes (CD45^+^ B220^+^) showed significant change upon the reconstitution of IL20RA (**Fig. 3, D and E**), which was consistent with the fact that IL20RA was screened in immunodeficient NOD-SCID mice lacking both T and B lymphocytes and natural killer (NK) cells (**Fig. 1A**).

To further confirm that IL20RA in OC cells affects the macrophages during the transcoelomic metastasis, we examined the immune cells in malignant ascites caused by the inoculation of SK-OV-3 cells with silenced IL20RA into NOD-SCID mice. FACS results show that the population of M1-like (MHCⅡ^+^) macrophages is significantly decreased and M2-like (CD206^+^) macrophages is significantly increased upon silencing IL20RA (**Fig. 3F**). Collectively, these data demonstrate that IL20RA in OC cells is able to regulate the polarization of peritoneal macrophages, instead of T lymphocytes and B lymphocytes, to prevent the transcoelomic spreading of OC cells into the peritoneal cavity.

### IL20RA mediates a direct conversation between OC cells and the macrophages that regulates the polarization of macrophages

To investigate if IL20RA supports a metastasis-preventing peritoneal immune-microenvironment through a direct crosstalk between OC cells and macrophages, we checked the polarization of macrophages *in vitro* under the stimulation of conditioned medium (CM) from IL-20-stimulated OC cells (**Fig. 4A**). CM from IL20RA-reconstituted ID8 cells (hereafter referred to as ‘CM^IL20RA^’) significantly stimulates the expression of M1-like markers while decreases the expression of M2-like markers in RAW 264.7 cells when compared with CM from empty vector-transfected ID8 control cells (referred to as CM^Vec^) (**Fig. 4B**). To further confirm, we prepared bone marrow-derived macrophages (BMDM) by treating the bone marrow cells (BMC) isolated from healthy C57BL/6 mice with 20 ng/mL macrophage colony stimulating factor (M-CSF) for 7 days, which were characterized by the adherent morphology and the expression of *Cd68* (**Fig. 4, C and D**). Consistently, CM^IL20RA^ dramatically induces the expression of M1-like markers and decreases the M2-like markers in BMDM (**Fig. 4E**).

**Figure 4.**
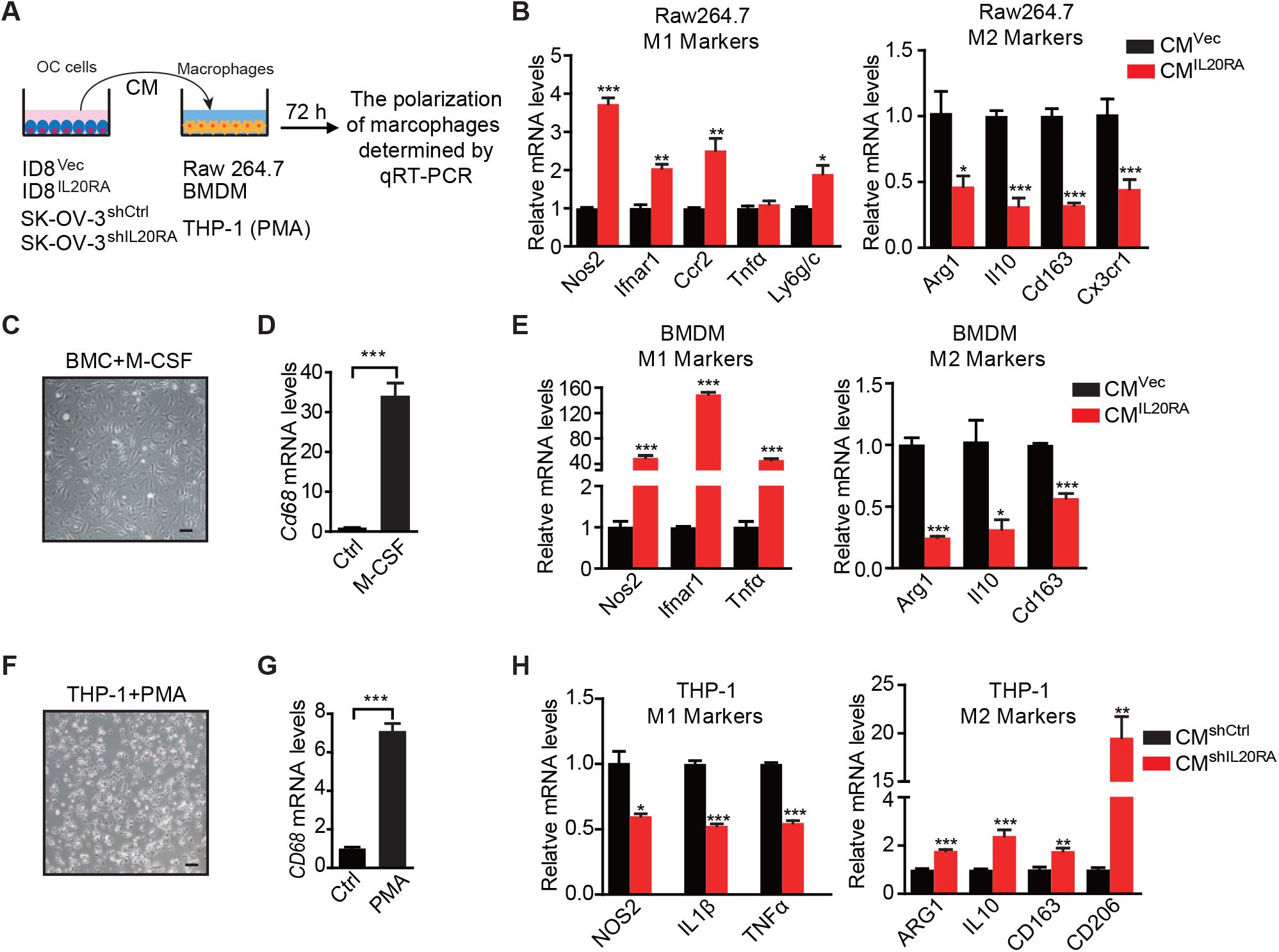
IL20RA mediates a direct crosstalk between OC cells and macrophages to regulate the polarization of macrophages. **A**. Schematic of the *in vitro* crosstalk experiment using conditioned medium (CM) from OC cells to educate macrophages. **B**. qRT-PCR analysis of macrophage marker genes in RAW 264.7 cells stimulated with CM of IL20RA-reconsitituted (CM^IL20RA^) or control (CM^Vec^) ID8 cells for 72 h. **C-D**. Representative image of BMDM differentiated from BMC with the treatment of M-CSF for 7 days (**C**), which were further characterized by qRT-PCR analysis of its marker gene *Cd68* (**D**). Scale bar: 20 μm. **E**. qRT-PCR analysis of macrophage marker genes in BMDM stimulated with CM^IL20RA^ or CM^Vec^ from ID8 cells for 72 h. **F-G**. Image of macrophages differentiated from THP-1 cells with the treatment of PMA for 48 h (**F**) and further characterization by qRT-PCR analysis of *CD68* (**G**). Scale bar: 20 μm. **H**. qRT-PCR analysis of macrophage marker genes in THP-1-derived macrophages stimulated with CM^shIL20RA^ or CM^shCtrl^ from shIL20RA or control shRNA (shCtrl) transfected SK-OV-3 cells for 72 h. All the qRT-PCR data are shown as means ± SEM from three independent experiments, *P < 0.05, **P < 0.01, ***P < 0.001, by unpaired two-sided student’s t-test.

In human macrophages differentiated from THP-1 monocytes by phorbol-12-myristate-13-acetate (PMA) treatment (**Fig. 4, F and G**), CM from IL20-stimulated, IL20RA-silenced SK-OV-3 cells significantly increases the expression of M2-like markers and decreases the expression of M1-like markers (**Fig. 4H**).

We also checked the polarization of macrophages *in vitro* using the indirect co-culture system (**Fig. S3E**). IL-20-stimulated, IL20RA-reconstituted ID8 cells have no effect on the migration capacity of RAW 264.7 cells (**Fig. S3, F and G**). However, co-culture with IL20RA-reconstituted ID8 cells dramatically induces the expression of M1-like markers and reduces the M2-like markers in RAW 264.7 cells (**Fig. S3H**). Besides, co-culture with IL-20-stimulated, IL20RA-silenced SK-OV-3 cells significantly increases the expression of M2-like markers and decreases the expression of M1-like markers in THP-1-derived macrophages (**Fig. S3I**). We also excluded the possibility that IL-20 directly regulated the polarization of macrophages since it did not affect the expression of both the M1-and M2-like markers in RAW 264.7 cells (**Fig. S3J**). Collectively, these data reveal a conserved, IL20RA-mediated direct crosstalk between OC cells and macrophages that favors an inflammatory immune-microenvironment.

### IL20RA-mediated education of macrophages plays essential roles in the prevention of the transcoelomic metastasis of OC

To investigate the function of the macrophages educated by IL20RA-mediated crosstalk with OC cells in the transcoelomic metastasis of OC, we injected ID8 cells alone, or together with CM^Vec^- or CM^IL20RA^-educated BMDM into the peritoneal cavity of C57BL/6 mice (**Fig. 5A**). CM^IL20RA^-educated BMDM significantly reduces the volume of malignant ascites (**Fig. 5, B and C**), and dramatically inhibits the formation of metastatic nodules in peritoneal cavity (**Fig. 5, D and E**), indicating that IL20RA-mediated crosstalk between OC cells and macrophages prevents the transcoelomic metastasis of OC. In addition, in peritoneal macrophage-depleted mice, reconstitution of IL20RA in ID8 cells can no longer reduce the volume of malignant ascites and inhibit the peritoneal metastasis of OC, indicating that IL20RA-mediated education of macrophages plays an essential role to suppress the transcoelomic metastasis of OC (**Fig. 5, F-J**).

**Figure 5.**
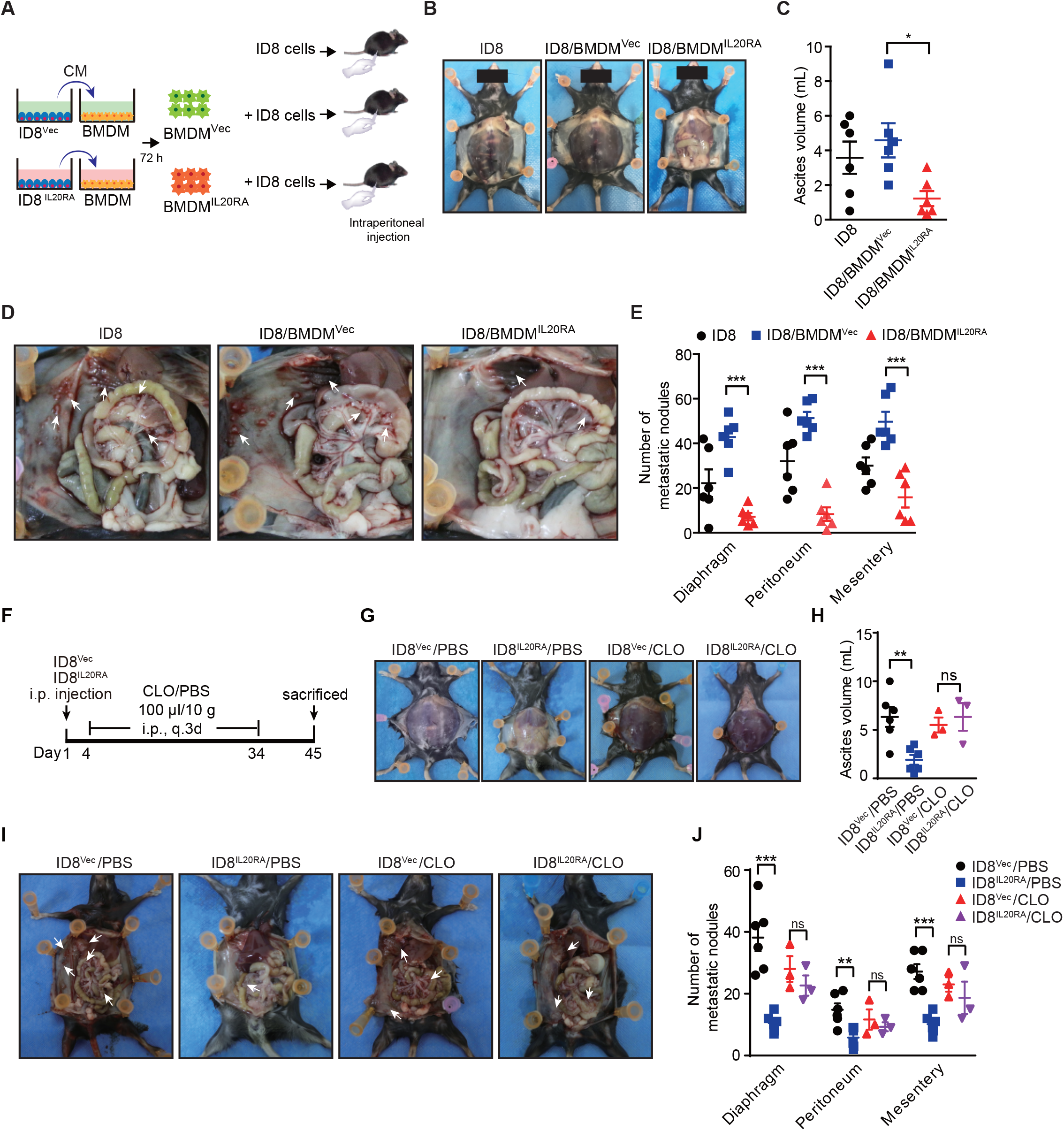
Macrophages play a prominent role in IL20RA-mediated suppression of OC metastasis. **A**. Schematic of the experiments. ID8 cells alone, or mixed with CM^IL20RA^- or CM^Vec-^stimulated BMDM, were injected into the peritoneal cavity of C57BL/6 mice. **B-E**. Representative images of the ascites formation (**B**) and metastatic nodules in peritoneal cavity (**D**) at day 45 post-inoculation. The quantification of ascites and metastatic nodules were shown in (**C**) and (**E**), respectively. Data are shown as means ± SEM, n = 6, *P < 0.05; **P < 0.01; ***P < 0.001, by unpaired two-sided student’s t-test. **F**. Schematic of the experiments. IL20RA-reconsitituted or control (Vec) ID8 cells were injected into the peritoneal cavity of C57BL/6 mice. The PBS liposomes (PBS) or clodronate liposomes (CLO) were intraperitoneal (i.p.) injected every 3 days. **G-J**. Representative images of the ascites formation (**G**) and metastatic nodules in peritoneal cavity (**I**) at day 45 post-inoculation. The quantification of ascites and metastatic nodules were shown in (**H**) and (**J**), respectively. Data are shown as means ± SEM, n = 6 (ID8^Vec^/PBS and ID8^IL20RA^/PBS), n = 3 (ID8^Vec^/C-LO and ID8^IL20RA^/CLO), ns not significant; **P < 0.01; ***P < 0.001, by unpaired two-sided student’s t-test.

### Mesothelial cells in peritoneum produce IL20RA ligands IL-20 and IL-24 when challenged with OC cells in the peritoneal cavity

To identify the signal sources in the peritoneal cavity that response to the disseminated OC cells through IL20RA, we injected ID8 cells into the peritoneal cavity of C57BL/6 mice to mimic the peritoneal dissemination of OC cells and checked the expression of possible IL20RA ligands, including IL-19, IL-20 and IL-24, in different tissues in the peritoneal cavity. A dramatic increase of *Il20* and *Il24* from the abdominal wall occurs upon the injection of OC cells into the peritoneal cavity (**Fig. 6A**), while there is no induction of these ligands in the intestine, ovary and peritoneal macrophages (CD11b^+^ F4/80^+^) (**Fig. 6, A and B**). In addition, reconstitution of IL20RA in ID8 cells does not induce these ligands in themselves (**Fig. 6C**). To further confirm, we isolated abdominal wall and co-cultured with ID8 cells *in vitro* and a boosted expression of *Il20* and *Il24* was observed (**Fig. 6D**). Hematoxylin and eosin (H&E) and IHC staining of the abdominal walls from mice bearing peritoneal-disseminated OC cells show that IL-20 and IL-24 are originated from mesothelial cells (**Fig. 6, E and F**), which are further confirmed by immunofluorescent (IF) staining (**Fig. S4, A and B**).

**Figure 6.**
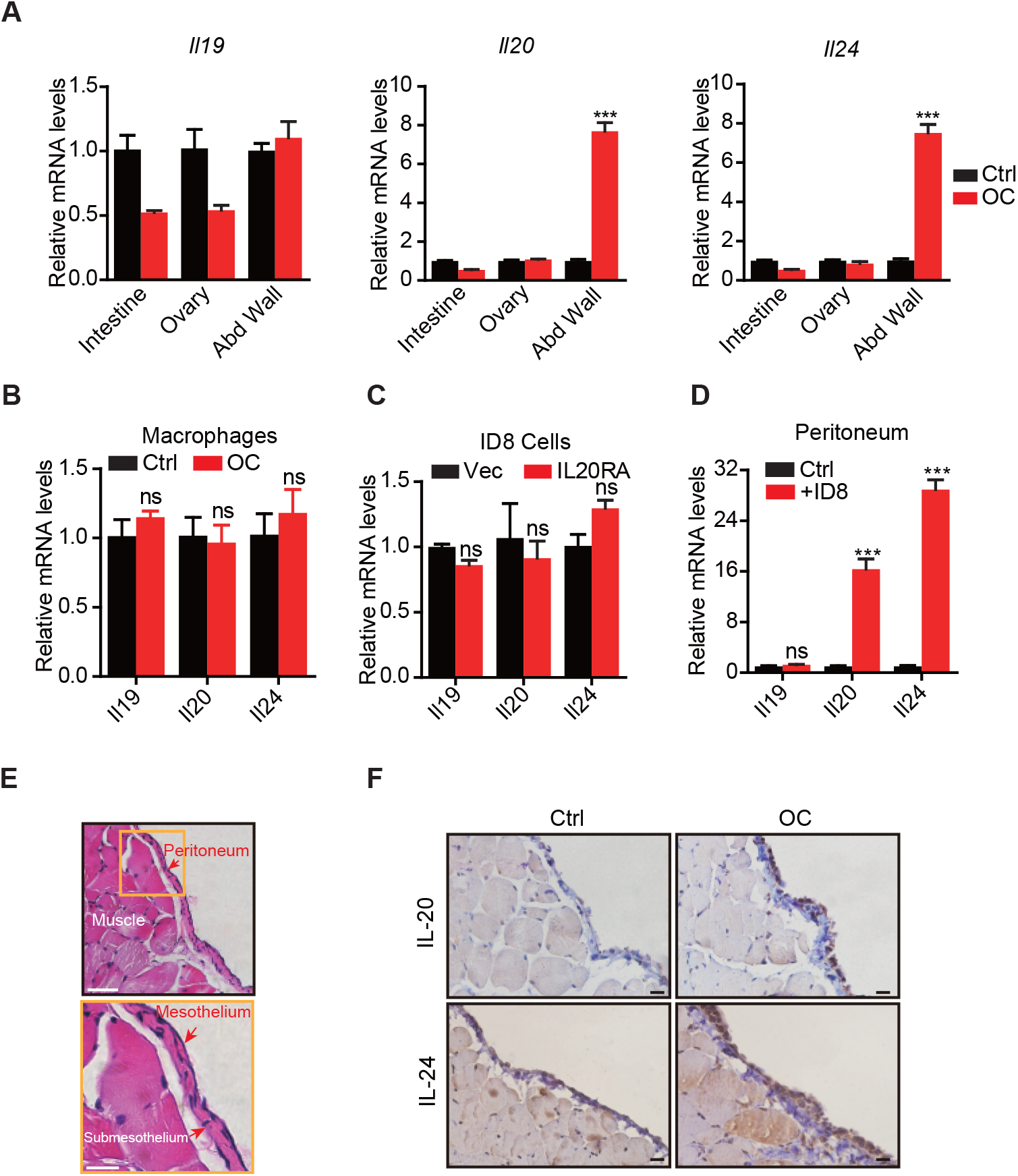
The mesothelial cells in peritoneum produce IL-20 and IL-24 when challenged by disseminated OC cells in the peritoneal cavity. **A-B**. qRT-PCR analysis of IL20RA ligands (*Il19, Il20* and *Il24*) in peritoneal organs (intestinal, abdominal wall, ovary) (**A**) or peritoneal macrophages (CD11b^+^ F4/80^+^) (**B**) taken from C57BL/6 mice with intraperitoneal injection of ID8 cells (OC) or PBS control (Ctrl) 9 days before. Data are shown as means ± SEM, n = 18 for Ctrl group and n = 6 for OC group, ***P < 0.001, ns not significant, by unpaired two-sided student’s t-test. **C**. qRT-PCR analysis of IL20RA ligands in IL20RA-reconstituted or control (Vec) ID8 cells (means ± SEM, ns not significant). **D**. qRT-PCR analysis of the abdominal walls dissected from C57BL/6 mice and co-cultured with medium (Ctrl) or ID8 cells for 48 h (means ± SEM from three independent experiments, ***P < 0.001, ns not significant, by unpaired two-sided student’s t-test). **E**. H&E staining of the abdominal wall of C57BL/6 mice. Scale bar: 50 μm (upper panel); 20 μm (lower panel). **F**. IHC staining of IL-20 and IL-24 in abdominal walls dissected from mice with intraperitoneal injection of ID8 cells (OC) or PBS control (Ctrl) 9 days before. Scale bar: 20 μm.

### IL-20/IL20RA activates OAS/RNase L-mediated NLR signaling to produce mature IL-18 for macrophage polarization

To explore the IL-20/IL20RA-mediated downstream signaling that modulates the polarization of peritoneal macrophage, we analyzed the transcriptome of ID8 cells with reconstituted IL20RA under the stimulation of IL-20. Kyoto Encyclopedia of Genes and Genomes (KEGG) enrichment analysis shows that the differentially expressed genes are enriched in the immune system (**Fig. S5A**). In particular, IL-20/IL20RA-mediated signaling greatly increases the expression of several genes, i.e., 2’-5’-oligoadenylate synthetase (*Oas1a* and *Oas1g*) and *Il18*, involved in OAS-RNase L-mediated NOD-like receptor (NLR) signaling pathway (**Fig. S5, B and C**), which regulates inflammasome signaling to activate Caspase-1 and hence the production of functional inflammatory cytokine IL-1β and IL-18 by cleavage during viral infections(Banerjee, 2016; Chakrabarti et al., 2015).

The induction of *Oas1a* and *Il18* in IL20RA-reconsitituted ID8 cells upon IL-20 stimulation is further confirmed by qRT-PCR (**Fig. 7A**) and enzyme linked immunosorbent assay (ELISA) for secreted IL-18 (**Fig. 7B**). Western blot analysis shows that IL-20/IL20RA triggers the phosphorylation of STAT3 and results in increased OAS1A, activated Caspase-1 and IL-18 by cleavage (**Fig. 7C**), which also occurs in human SK-OV-3 cells when stimulated by IL-20 (**Fig. 7, D and E**). In addition, in syngeneic murine OC model, IL20RA-reconstituted ID8 cells in intraperitoneal cavity causes significantly higher amount of IL-18 in ascites when compared with empty vector transfected cells (**Fig. 7F**). We further identify a STAT3-binding site on the promoter of *Oas1a* gene by chromatin immunoprecipitation and qPCR (ChIP-qPCR) (**Fig. 7, G and H**), confirming that IL-20/IL20RA induces *Oas1a* through downstream STAT3. Consistently, IHC staining of human OC tissues shows that the levels of phosphorylated STAT3, OAS1, and IL-18 are significantly lower in metastatic lesions than those in primary sites (**Fig. 7, I and J**). In addition, transcriptome analysis on a cohort of 530 serous OC patients from TCGA database shows the positive correlation of *IL20RA* with both *OAS1* and *IL-18* (**Fig. S5D**), further supporting IL-20/IL20RA activates OAS/RNase L-mediated inflammasome signaling in human OC as well.

**Figure 7.**
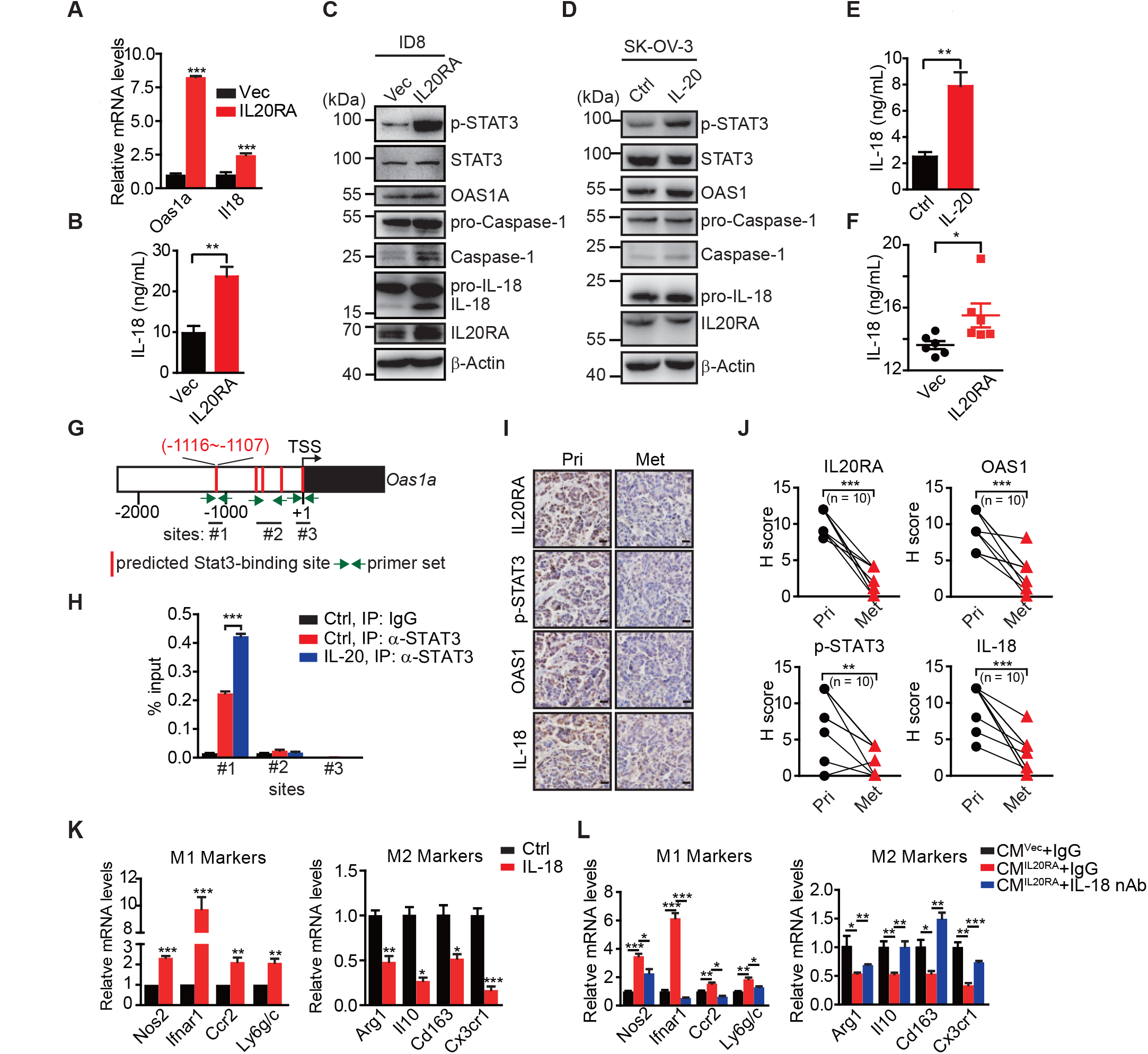
IL-20 activates the IL20RA-STAT3-OAS1/RNase L-NLR signaling to produce IL-18 to regulate the macrophages polarization. **A.** qRT-PCR analysis of *Oas1a* and *Il18* upon IL20RA reconstitution in ID8 cells. **B**. The secreted IL-18 from IL20RA-reconstituted and control ID8 cells was measured by ELISA. **C.** Western blot analysis of indicated proteins in IL20RA-reconstituted and control ID8 cells. **D**. Western blot analysis of indicated proteins in SK-OV-3 cells stimulated with IL-20 or PBS (Ctrl) for 24 h. **E**. ELISA measurement of IL-18 secreted from SK-OV-3 cells stimulated by IL-20 for 24 h. **F**. ELISA measurement of IL-18 in ascites from mice with intraperitoneal injection of IL20RA-reconstituted or control (Vec) ID8 cells 40 days post-injection (n = 6 for each group). **G**. Schematic of the *Oas1a* promoter with predicted STAT3-binding sites and the primer sets. **H**. IL20RA-reconstituted ID8 cells were stimulated with IL-20 or PBS (Ctrl) for 24 h before the STAT3 binding on *Oas1a* promotor was analyzed by ChIP-qPCR. **I-J**. IHC analysis of IL20RA, p-STAT3, OAS1 and IL-18 in human primary OC tissues (Pri) and paired peritoneal metastatic nodules (Met) (**I**) and quantification (**J**). **P < 0.01; ***P < 0.001 by paired two-sided student’s t-test. Scale bar: 20 μm. **K**. qRT-PCR analysis of macrophage marker genes in RAW 264.7 cells stimulated with IL-20 protein for 72 h. **L**. qRT-PCR analysis of macrophage marker genes in RAW 264.7 cells treated by CM^Vec^ or CM^IL20RA^ together with IL-18 neutralizing antibody (nAb) or nonspecific lgG (IgG) for 72 h. All the qRT-PCR and ELISA data are shown as means ± SEM from three independent experiments, *P < 0.05, **P < 0.01, ***P < 0.001, by unpaired two-sided student’s t-test.

To get insights into the role of IL-18 in macrophage polarization, RAW 264.7 cells were treated with IL-18, which resulted in greatly increased expression of M1-like markers and significantly decreased M2-like markers (**Fig. 7K)**. CM^IL20RA^-induced polarization of RAW 264.7 cells to M1-like phenotypes can be dramatically blocked by IL-18 neutralization antibody (**Fig. 7L**), highlighting IL-18 as the essential factor downstream IL-20/IL20RA signaling to regulate the polarization of macrophages.

### Administration of IL-18 protein strongly suppresses the transcoelomic metastasis of OC

Given the dramatic decrease of IL20RA in the peritoneal metastasized OC cells (**Fig. 2**) and the essential role of IL-18 as a major IL20RA downstream factor inhibiting peritoneal dissemination, we postulated that the direct administration of IL-18 might be an easy strategy to suppress the peritoneal growth of OC. To test, we treated the mice bearing orthotopically transplanted ID8 xenografts with recombinant IL-18 protein (**Fig. 8A**). It shows that intraperitoneal injection of IL-18 dramatically reduces the ascites formation and the numbers of metastatic nodules in diaphragm, peritoneum and mesentery (**Fig. 8, B-E**). Furthermore, direct administration of IL-18 significantly increases the proportion of M1-like (MHCⅡ^+^ CD206^−^) macrophages and decreases M2-like (MHCⅡ^−^ CD206^+^) macrophages in the malignant ascites (**Fig. 8F**).

**Figure 8.**
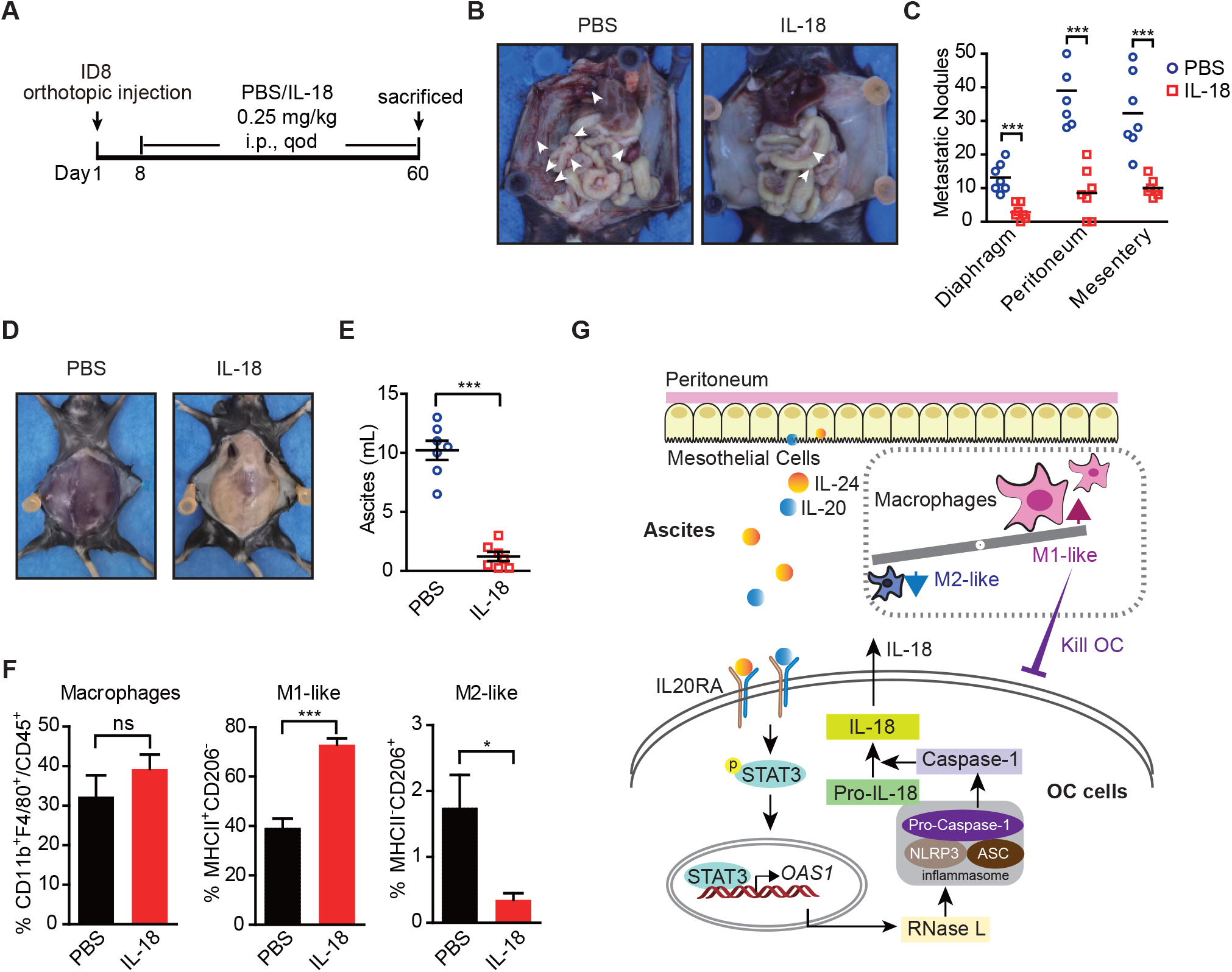
The therapeutic effect of recombinant IL18 against the metastasis of OC. **A**. Schematic of the experiments. ID8 cells were orthotopic injected into the ovaries of C57BL/6 mice. The PBS or IL-18 protein were intraperitoneal (i.p.) injected every 2 days. **B-E**. Representative images of the metastatic nodules in peritoneal cavity (**B**) and ascites formation (**D**) at day 60 post-inoculation. The quantification of metastatic nodules and ascites were shown in (**C**) and (**E**), respectively. Data are shown as means ± SEM (n=7). ***P < 0.001, by unpaired two-sided student’s t-test. **F**. Flow cytometry analysis of macrophages (CD45^+^ CD11b^+^ F4/80^+^) and M1-like (MHCⅡ^+^ CD206^−^) and M2-like (MHCⅡ^−^ CD206^+^) subpopulations in ascites formed in C57BL/6 mice at 60 days after orthotopically inoculated with ID8 cells (means ± SEM, n = 5, *P < 0.05, ***P < 0.001, ns not significant, by unpaired two-sided student’s t-test). **G**. Schematics summarizing the IL-20/IL20RA-OAS1/RNase L-NLR-IL-18 axis in preventing the transcoelomic metastasis of OC.

In conclusion, we discovered a new crosstalk of disseminated OC cells with peritoneal mesothelial cells and macrophages through IL-20/IL20RA/IL-18 axis in shaping the peritoneal immunomicroenvironment to prevent the transcoelomic metastasis of OC cells. OC cells, when disseminates into the peritoneal cavity, stimulate the mesothelial cells of the peritoneum to produce IL-20 and IL-24, which in turn activate the IL20RA downstream signaling in OC cells to trigger the OAS/RNase L-mediated NLR inflammasome signaling to produce mature IL-18, which consequently promotes the polarization of peritoneal macrophages into M1-like subtype to clear invaded OC cells in peritoneal cavity (**Fig. 8G**). Highly metastatic OC cells always block this pathway by decreasing IL20RA expression, suggesting that reactivation its downstream signaling, such as the application of IL-18, could be a useful strategy for the therapy of OC at advanced stages.

## Discussion

The common metastasis mode of OC is transcoelomic metastasis, in which the cancer cells disseminate into the peritoneal cavity and colonize on the peritoneum and abdominal organs. Transcoelomic metastasis occurs in about 70% of OC patients and many other types of cancers, such as pancreatic cancer, gastric cancer, endometrial cancer and colon cancer(Mikula-Pietrasik et al., 2018). Patients with transcoelomic metastasis are diagnosed at stage III and IV, which means high mortality and poor prognosis. The mechanisms, in particular, the peritoneal specific factors, behind this process still remain largely unclear. Here, through a genome-scale gene knockout screening in the orthotopic murine model of OC, we reveal an essential role of IL-20/IL20RA-mediated crosstalk between cancer cells and peritoneum mesothelial cells in regulating the polarization of peritoneal macrophages to prevent transcoelomic metastasis. This peritoneal-specific immune modulation mechanism may also happen in other types of cancers that can disseminate into the peritoneal cavity, such as gastric cancer, endometrial carcinoma, colon cancer and bladder carcinoma, in which the high expression of IL20RA also correlates with a better clinical outcome of patients (**Fig. S6**), highlighting the importance of IL-20/IL20RA-mediated epithelial immunity in preventing cancer metastasis into the peritoneal environment.

Immune cell-produced IL-20 subfamily cytokines and their receptors mainly expressed in epithelial cells facilitate the crosstalk between leukocytes and epithelial cells, which has essential functions in epithelial innate immunity for the host defence and tissue repair at epithelial surfaces. Due to the complicated effects of IL-20 subfamily cytokines in inflammation stimulation upon infection or injury and later immunosuppression for wound repair and tissue homeostasis restoration, both tumor-promoting and tumor-suppressing roles of IL-20 subfamily cytokines have been reported. IL-20 and IL-26 were discovered to promote proliferation and migration of bladder cancer, breast cancer and gastric cancer cells, while IL-24 was found to inhibit the proliferation and metastasis of melanoma, prostate cancer and ovarian cancer(Fisher et al., 2007; Gopalan et al., 2007; Hsu et al., 2012; Lee et al., 2013; Pradhan et al., 2018; You et al., 2013). Although IL-20 subfamily cytokines have been extensively reported to activate STAT3 to exhibit oncogenic effects, STAT3 has also been shown to be able to switch its roles from tumor-promoting at early stages to tumor-suppressing in the invasion process(Musteanu et al., 2010). The different roles of IL-20 subfamily cytokines in tumor growth and metastasis may depend on the local inflammatory environment that involves a complex network of conversations between cancer cells and various stromal components. For OC metastasized to omentum, cancer cell-mediated conversion of omentum stromal cells, including adipocytes, fibroblasts, macrophages and mesenchymal stem cells, to cancer-associated stromal cells is of vital importance for the metastatic growth of OC cells on omentum (Motohara et al., 2019; Thibault et al., 2014). The abdominal cavity is lined with a layer of mesothelial cells that form the first line of defence against microbial pathogens through the expression of retinoic acid-inducible gene-I-like (RIG-I–like) receptors, Toll-like receptors and C-type lectin-like receptors(Colmont et al., 2011; Kato et al., 2004; Park et al., 2007). Besides, mesothelial cells can secret cytokines, chemokines and growth factors to regulate the inflammatory responses in peritoneal cavity(Isaza-Restrepo et al., 2018; Yao et al., 2004). However, whether and how mesothelial cells response to cancer cells invaded into the peritoneal cavity remain largely unknown. It is more difficult for OC cells to attach to mesothelial cells compared with the extracellular matrix and the fibroblasts(Aziz et al., 2019; Kenny et al., 2007; Niedbala et al., 1985). During the metastatic growth, OC cells can destroy the mesothelial cell layer by promoting the mesothelial-to-mesenchymal transition (MMT) of mesothelial cells and hence the loss of cell-cell adherence(Iwanicki et al., 2011; Kenny et al., 2011; Sandoval et al., 2013). Here, we revealed, for the first time, that mesothelial cells can actively respond to OC cells invaded into the peritoneal cavity by secreting IL-20 and IL-24 to communicate with OC cells, consequently resulting in a further crosstalk between OC cells and macrophages for the formation of an anti-tumor microenvironment. This mesothelial-initiated defence mechanism is often times overcome by successfully metastasized OC cells through the downregulation of IL-20/IL-24 receptor IL20RA (**Fig. 2, A-F**) to shut down the communication between OC and mesothelial cells.

It has been reported that in OC patients, the majority of cell types in ascites are lymphocytes (37%) and macrophages (32%), contributing to the progression and metastasis of OC(Kipps et al., 2013). Generally, macrophages have the potentials to differentiate into cancer-inhibiting M1-subtype and cancer-promoting M2-subtype. The ratio of M1/M2-subtype macrophages has predictive value for the prognosis of OC patients, with a higher M1/M2-subtype ratio being associated with a better prognosis, while a higher CD163^+^ (M2-subtype)/CD68^+^ (total macrophages) ratio with a poor prognosis(Reinartz et al., 2014; Zhang et al., 2014). Re-polarization of macrophages to increase the ratio of M1/M2 macrophages in ascites is a promising strategy for the treatment of OC. It has reported that host-produced histidine-rich glycoprotein (HRG) could inhibit the growth and metastasis of tumors *via* re-polarizing the macrophages from M2-subtype to M1-subtype(Rolny et al., 2011). Nanoparticles encapsulating miR-125b could specifically re-polarize the peritoneal macrophages to M1-subtype, thereby improving the efficacy of paclitaxel in OC(Parayath et al., 2019). Here, we reveal IL20RA as a key receptor in regulating the polarization of peritoneal macrophages, which is silenced in disseminated OC cells for the accumulation of M2-subtype macrophages in the peritoneal cavity for a successful metastatic growth. This indicates that reactivation of IL-20/IL20RA signaling by the application of IL-20 or IL-24 may not be a useful strategy to prevent OC transcoelomic metastasis. However, we also identify the key downstream effector of IL-20/IL20RA signaling that promotes M1-like macrophage, IL-18, which is usually involved in the regulation of innate immune responses against various pathogens (Fabbi et al., 2015).

IL-18 is an immunoregulatory cytokine with anti-cancer effect in many types of cancers(Coughlin et al., 1998; Nishio et al., 2008; Salcedo et al., 2010; Zaki et al., 2010). The normal ovarian and colon epithelia have the capacity to process and release active IL-18, thereby supporting a local anti-tumor immune microenvironment. However, during the malignant transformation, the ovarian and colon cancer cells lose the ability to process IL-18 as a result of the defective Caspase-1 activation(Feng et al., 2005; Wang et al., 2002). Thus, OC cells express more pro-IL-18 but poorly produce mature-IL-18. Elevated IL-18 was reported in the ascites and sera of OC patients in a 23-kDa pre-form, whereas the 18-kDa mature and active form was undetectable (Orengo et al., 2011). Here, we show that reconstitution of IL20RA in highly metastatic OC cells, can reactivate the production of mature IL-18 *via* IL-20/IL20RA-OAS1/RNase L-NLR axis, suggesting that the silence of IL20RA to be a possible reason for the loss of mature IL-18 in OC. IL-18 has been shown to exert antitumor activity through the activation of T cell(Fabbi et al., 2015). Here, our results expand its roles to promote the polarization of macrophages to inflammatory M1-like subtype to exhibit a potent anti-tumor effect against OC. Several clinical trials testing the anti-tumor efficacy of IL-18 have been conducted in lymphomas and melanoma(Robertson et al., 2013; Robertson et al., 2006; Tarhini et al., 2009). Our study shows that the administration of recombinant IL-18 significantly suppresses the metastasis of IL20RA-deficient OC cells, suggesting the great value of IL-18 in the therapy of OC at advanced stages.

## Materials and Methods

### Cell culture and cytokine treatment

THP-1, RAW 264.7, A2780 and HEK 293T cells were purchased from the American Type Culture Collection (ATCC, Washington, USA). SK-OV-3 and ES-2 cells were purchased from the Cell Bank of the Chinese Academy of Sciences (Shanghai, China). ID8 cells were purchased from Merck (Darmstadt, Germany) and screened in C57BL/6 mice for highly metastatic and HGSOC-like ID8 cells as described by Kristy K Ward and colleagues(Diaz Osterman et al., 2019; Ward et al., 2013). SK-OV-3 and OVCAR-5 cells were cultured in McCoy’s 5A Medium Modified (Biological Industries, Israel). ES-2, A2780, ID8 and HEK 293T cells were cultured in Dulbecco’s modified Eagle’s medium (DMEM) with high glucose (Biological Industries). THP-1 and RAW 264.7 cells were cultured in RPMI-1640 (Biological Industries). All the mediums were supplemented with 10% fetal bovine serum (Biological Industries), 100 U/mL penicillin and 100 μg/mL streptomycin (Gibco, Grand Island, NY, USA) and maintained at an atmosphere of 5% CO2 and 95% air at 37 °C.

To study the role of IL-18 and IL-20 protein in the polarization of RAW 264.7 cells, RAW 264.7 cells were treated with 5 ng/ml murine recombinant IL-18 protein or 5 ng/mL murine recombinant IL-20 protein (Origene, Rockville, MD, USA) for 72 hours, respectively. To detect the activation of signaling pathway, ID8 and SK-OV-3 cells were stimulated with 2.5 ng/mL murine and human recombinant IL-20 protein (Sino Biological, Beijing, China) for 24 hours, respectively.

### The differentiation of THP-1 and bone marrow cells (BMCs)

THP-1 was differentiated into macrophages by the treatment with 100 ng/mL PMA (Sigma-Aldrich, St Louis, MO, USA) for 48 hours. BMCs were isolated from the femurs and tibias of C57BL/6 mice. The bone marrow in the femurs and tibias were flushed and collected with DMEM/F-12. Erythrocytes in bone marrow were lysed using Red Blood Cell Lysis Buffer (Solarbio, Beijing, China). BMCs were cultured in DMEM/F-12 supplemented with 20 ng/mL murine M-CSF (315-02-10, Peprotech, Cranbury, NJ, USA) in a density of 4 × 10^6^/mL. The medium was changed every 3 days. BMCs were differentiated into BMDM with the stimulation of M-CSF for 7 days. 5 mM of EDTA was used to detach the cells from the culture dishes. The detached cells were seeded in the six-well plates for subsequent treatment with CM from cancer cells.

### Establishment of stable cell lines

Knocking down IL20RA was achieved by the insertion of shRNA templates into the lentivirus-based RNAi vector pLV-H1-EF1α-puro (Biosettia, San Diego, CA, USA). The sequence of shRNA targeting human *IL20RA* (shIL20RA) is 5’ AAAAGCCCGCAAACGTTACAGTACTTTGGATCCAAAGTACTGTAACGTTT GCGGGC 3’. The control shRNA plasmid targeting Escherichia coli *lacZ* gene (shCtrl) was purchased from Biosettia with the sequence of 5’ AAAAGCAGTTATCTGGAAGATCAGGTTGGATC AACCTGATCTTCCAGATAACTGC 3’. The coding sequence of mouse *Il20ra* was synthesized by Sangon Biotech (Shanghai, China) and cloned into plasmid pLV-EF1α-MCS-IRES-Bsd (Biosettia). All plasmids were validated by sequencing. The procedures of lentivirus package and infection were described previously(Chang et al., 2015). Cells infected with lentivirus for RNAi or reconstituted expression were selected with 2.5 μg/mL of puromycin and 2.5 μg/mL of blasticidin (Thermo-Fisher Scientific, Waltham, MA, USA), respectively.

### CRISPR/Cas9 knockout *in vivo* screening

GeCKO v2 human library made by Zhangfeng’s lab was purchased from Addgene (Watertown, MA, USA) amplified as described(Joung et al., 2017). The library contains 122 756 sgRNAs targeting 19 050 genes and 1 864 miRNAs, and 1 000 non-targeting sgRNAs. Lentivirus carrying the library were generated and infected SK-OV-3 cells with the multiplicity of infection (MOI) around 0.5. After the selection with puromycin, two million SK-OV-3 cells expressing sgRNAs were orthotopically injected into NOD-SCID mice. The mice were sacrificed after 40 days. Primary tumors were dissected. Metastatic nodules in the peritoneal cavity were isolated, *in vitro* expanded for next run of orthotopic injection. After 3 rounds of *in vivo* selection, peritoneal metastatic cells were collected for subsequent high throughput DNA deep sequencing to identify metastasis-related candidate genes. The RIGER P analysis was used to analyze the data of sgRNA libraries sequencing.

### Murine OC models

Six to eight weeks old female NOD-SCID and C57BL/6 mice were purchased from SPF Biotechnology (Beijing, China). Two million of human SK-OV-3 cells or five million of murine ID8 cells suspended in 20 μL phosphate-buffered saline (PBS) were orthotopically injected into the left ovary of NOD-SCID and C57BL/6 mice, respectively. Ascites volumes, the numbers of metastatic nodules were measured and total cells in ascites were harvested 40 days post-inoculation for SK-OV-3 OC xenograft model and 60 days post-inoculation for ID8 OC syngeneic model. For the intraperitoneal OC mouse model, five million of ID8 cells suspended in 100 μL of PBS were injected directly into the peritoneal cavity of C57BL/6 mice. At day 45 post-injection, ascites and the number of metastatic nodules were analyzed.

To investigate the role of peritoneal macrophage in IL20RA-mediated OC metastasis, five million of ID8 cells alone or together with 1.25 × 10^6^ of BMDM (educated by CM from ID8 cells for 72 hours) in 100 μL of PBS were directly injected into the peritoneal cavity of C57BL/6 mice. In another model, five million of IL20RA-reconstituted or control ID8 cells were injected into the peritoneal cavity of C57BL/6 mice. The mice were then intraperitoneal injected with PBS liposomes or clodronate liposomes (100 μL/10 g of mouse weight) (CP-030030, LIPOSOMA, Netherlands) every 3 days from day 4 to day 34 post-injection.

To identify the source of IL20RA ligands in peritoneal cavity, five million of ID8 cells in 100 μL PBS (OC) or only 100 μL PBS (control) were directly injected into the peritoneal cavity of C57BL/6 mice. The mice were sacrificed 9 days post-injection. Peritoneal flushing fluid by PBS were collected. Total cells in peritoneal cavity were harvested for sorting the peritoneal macrophages. The total RNAs from intestine, abdominal wall and ovary were extracted by using TRIzol reagent (Thermo-Fisher Scientific) for subsequent gene expression analysis.

To investigate the therapeutic effect of IL-18, five million of murine ID8 cells suspended in 20 μL PBS were orthotopically injected into the left ovaries of C57BL/6 mice. The mice were intraperitoneal injected with PBS or IL-18 protein (0.25 mg/kg of mouse weight) every 2 days from day 8 to day 60 post-injection.

### Analysis of OC patient samples

OC specimen from human patients were obtained from Tianjin Center Hospital of Gynecology Obstetrics with informed consent. Twenty fresh paired samples, including primary serous OC tissues, ascites and metastasis nodules in peritoneal cavity, were obtained from OC patients during operation between 2017 and 2019. The inclusion criteria included: serous ovarian adenocarcinoma, age > 40 and TNM stage Ⅲ - Ⅳ. Primary tumor tissues and metastases samples were dissected into two parts: one stored in 4% paraformaldehyde for IHC staining, the other stored in TRIzol reagent for the extraction of protein and RNA. Cancer cells in ascites were harvested for the extraction of protein and RNA. The histopathology and TNM stages of OC patients were confirmed by pathologist and clinical doctors.

The analysis on the correlation between the prognosis of OC patients and the expression of IL20RA or IL20RB were performed using the online Kaplan Meier plotter tool (http://www.kmplot.com/) on serous OC cohorts (Gyorffy et al., 2012). The co-expression analysis on IL20RA with IL18 and OAS1 in OC patients were examined using the data from TCGA database (Ovarian Serous Cystadenocarcinoma, Provisional, n = 530). We used mRNA RNA-seq on pan-cancer section in the online Kaplan Meier plotter tool to analyze the correlation between the prognosis of other cancer types and the expression of IL20RA.

### Protein extraction and western blot

The tissues were homogenized by TissueLyser (SCIENTZ-48, Ningbo, Zhejiang, China). Total protein in tissues was extracted using TRIzol reagent. Cells were lysed in RIPA lysis buffer with protease inhibitor cocktail (Roche Life Science, Switzerland) and phosphatase inhibitor cocktail (Sigma-Aldrich). The concentrations of proteins were measured by Pierce^™^ BCA Protein Assay Kit (Thermo-Fisher Scientific), and 20 μg of total protein were loaded into Tris-acrylamide gels. Proteins were transferred onto PVDF membranes (Merck). The membranes were blocked with 5% defatted milk before blotted with primary antibodies and secondary antibodies [anti-mouse IgG-HRP (at 1:2,000 dilution), anti-rabbit IgG-HRP (at 1:2,000 dilution) (ZSGB-BIO, Beijing, China)]. The membranes were incubated with Immobilon® Western Chemiluminescent HRP Substrate (Merck) and visualized and photographed by Tanon-5200 (Tanon, Shanghai, China). The primary antibodies used in western blot are listed in Supplemental Table 1.

### IL-18 protein expression and purification

The coding sequence of the mature IL-18 was cloned into pET-20b plasmid (Promega) before being transduced into *E. coli* BL21 (DE3) cells (CWBIO, Beijing, China). The expression of recombinant IL-18 was induced by 0.5 mM IPTG (Thermo-Fisher Scientific) for 8 h at 37 °C. Cells were harvested and lysed by sonication in the lysis buffer (300mM NaCl, 50mM NaH2PO4 pH 8.0, 10mM imidazole and 2 mg/ml lysozyme). The supernatants were incubated with Ni-NTA agarose beads (Qiagen) overnight at 4 °C. The beads were consecutively washed with wash buffer containing 20 mM imidazole (300 mM NaCl, 50 mM NaH2PO4 pH 8.0, 20 mM imidazole). The recombinant proteins were eventually eluted with elution buffer containing 250 mM imidazole (300 mM NaCl, 50 mM NaH2PO4 pH 8.0, 250 mM imidazole) and were dialyzed to remove imidazole. The endotoxin in the purified protein was removed using High Performance Endotoxin Removal Agarose Resin (20518ES10, Yeasen Biotech, Shanghai, China) and examined by using Kinetic Turbidimetric LAL (60401ES06, Yeasen Biotech) (final concentration of endotoxin < 0.03 EU/mL).

### qRT-PCR

One microgram of total RNAs from cells and tissues were reversely transcribed into cDNAs using M-MLV reverse transcriptase (Promega, Madison, WI, USA). Quantitative PCR analysis was performed by using SYBR Green SuperMix (Yeasen Biotech) on LightCycler®96 system (Roche) with the program as follows: 95℃ for 300 s, 45 cycles of two-step reaction, i.e., 95 ℃ for 30 s followed by 60 ℃ for 45 s. The primer sequences are listed on Supplemental Table 2. The relative gene expression was calculated by the 2^−ΔΔCt^ method with β-actin used for the normalization.

### H&E staining, IHC and IF

The tissues dissected from patients and mice were fixed in 4% paraformaldehyde, dehydrated, paraffin-embedded, consecutive sectioned in 6-μm thickness, and then deparaffinized. For H&E staining, tissue sections were stained with hematoxylin and eosin (Origene). For IHC staining, after treated with antigen retrieval solution (Solarbio) and 3% hydrogen peroxide, tissue sections were blocked with 5% goat serum, incubated with primary antibodies, biotin-conjugated secondary antibodies (Vector Laboratories, Burlingame, CA, USA) and streptavidin-HRP (Vector Laboratories, Burlingame, CA, USA). The sections were visualized with 3,3’-diaminobenzidine (DAB) substrate (ZSGB-BIO, Beijing, China), then counter stained with hematoxylin. Images were taken using an Olympus BX53 microscope (Tokyo, Japan). H score was used to quantify the degree of immunostaining, which was calculated by multiplying positively stained area (P) with staining intensity (I). The degrees for P were 0 - 4 (0, < 5%;1, 5% ∼ 25%; 2, 25% ∼ 50%; 3, 50% ∼ 75%; 4, 75%∼100%). The degrees for I were 0 - 3 (0, none; 1, weak; 2, moderate; 3, strong). For IF staining, tissue sections were blocked with 5% goat serum, incubated with primary antibodies, secondary antibodies conjugated with Alexa Fluor-488 and Alexa Fluor-594 (Thermo-Fisher Scientific), then counter stained with 4’,6-diamidino-2-phenylindole (DAPI, Sigma-Aldrich). Images were taken using an Olympus FV1000 confocal microscope (Tokyo, Japan). The antibodies used in IHC and IF staining are listed in Supplemental Table 1.

### Flow cytometry

Total ascites was collected when the mice were sacrificed. Erythrocytes in ascites were lysed using Red Blood Cells Lysis Buffer (Solarbio). Cells were washing using PBS with 1% FBS, and passed through a 40 μm nylon filter (BD Biosciences, San Diego, CA, USA), then cells were Fc-blocked with anti-CD16/32, and stained with CD45-PE, CD11b-APC, F4/80-APC/CY7 antibodies for detecting macrophages. MHC Ⅱ -PERCP/Cy5.5 and CD206-FITC antibodies were used to differentiate M1- and M2-subtype macrophages. For detecting T cells and B cells, cells were stained with CD45-PE, CD3-FITC, CD4-APC, CD8-PE/CY7, CD45R/B220-FITC antibodies. Cells were incubated with the antibodies for 30 min in the dark, and washed twice using PBS with 1% FBS. LSRFORTESSA (BD Biosciences, San Diego, CA, USA) was used to obtained the data and FlowJo software (BD Biosciences, San Diego, CA, USA) was used to analyze the data. The antibodies used in flow cytometry are listed on Supplemental Table 1.

### Isolation of peritoneal macrophages

Total cells in peritoneal cavity of healthy C57BL/6 mice were harvested by peritoneal lavage using PBS. As the staining protocol described in flow cytometry, cells from peritoneal flushing fluid of healthy mice and ascites of OC mice were stained with CD11b-APC and F4/80-APC/CY7 antibodies. CD11b^+^ F4/80^+^ peritoneal macrophages were isolated from total cells using FACSAria SORP (BD Biosciences, San Diego, CA, USA), and processed for RNA extraction.

### Cell proliferation and migration assay

The proliferation of ID8 and SK-OV-3 cells were examined by cell counting under the stimulation of IL-20. Two thousand of ID8 cells and SK-OV-3 cells were seeded in 96-well plates evenly. The cells were digested and counted by using the Invitrogen Countess II FL Automated Cell Counter (Thermo-Fisher Scientific) every 24 hours until they reached nearly 100% confluence.

Transwell Permeable Supports System (Corning Inc., Corning, NY, USA) was used for SK-OV-3 cells migration assays. 2×10^4^ cells in 200 μl medium containing 2% FBS were seeded into the top chamber of an 8 μm Millipore transwell insert in a 24-well plate, and medium containing 10% FBS was added to the bottom chamber. After 8 hours, cells migrated to the bottom surface of the transwell membranes were fixed in 4% paraformaldehyde and stained with crystal violet (Beyotime, Shanghai, China). The migration ability of RAW 264.7 cells was examined using indirect transwell cell culture system (8 μm, Millipore). 4 × 10^5^ of ID8 cells were seeded in 6-well plates evenly. After 24 hours, 1 × 10^5^ of RAW 264.7 cells were seeded in the upper transwell chamber. After co-cultured for 24 hours, cells migrated to the bottom surface of the transwell membranes were fixed in 4% paraformaldehyde and stained with crystal violet. The membranes were washed by PBS. Images of the membranes were taken using an Olympus BX53 microscope (Tokyo, Japan), and the numbers of migrated cells were counted in five randomly chosen fields.

### CM stimulation and co-culture system

IL20RA-silenced SK-OV-3, IL20RA-reconstituted ID8 or their parallel control cells were cultured in 10-cm dishes for 72 hours with the stimulation of IL-20 protein, the CM of them were collected and centrifuged at 2,000 rpm for 10 minutes to obtain the supernatants. 2 × 10^5^ of RAW 264.7 cells, 1 × 10^6^ of BMDM and 1 × 10^6^ of THP-1-derived macrophages were seeded in 6-well plates 24 hours before the medium was changed to fresh medium mixed with OC cell-derived CM at the ratio of 2:1 and incubated for 72 hours. The macrophages were then collected in TRIzol reagent for RNA extraction. To investigate the role of IL-18 in regulating the polarization of macrophages, RAW 264.7 cells were stimulated with the ID8 cell-derived CM supplemented with 5 μg/mL of IL-18 neutralizing antibody (R&D, Minneapolis, MN, USA) or 5 μg/mL of rat IgG1 Isotype as a control (Thermo-Fisher Scientific).

Macrophages and OC cells were indirect co-cultured using 0.4 μm pore membrane transwell (Merck). 2 × 10^5^ of RAW 264.7 cells and 1 × 10^6^ of THP-1-derived macrophages were seeded in 6-well plates and cultured for 24 hours before 1 × 10^5^ of SK-OV-3 cells or 5 × 10^4^ of ID8 cells stimulated with IL-20 protein were seeded in the upper transwell chamber. After co-cultured for 72 hours, macrophages were collected in TRIzol reagent for subsequent gene expression analysis. To investigate whether OC cells induces the production of IL-20 and IL-24 from peritoneum *in vitro*, abdominal wall and OC cells were co-cultured. 2 × 10^5^ of ID8 cells were seeded in 6-well plates. After 24 hours, 2 × 2 cm^2^ abdominal wall tissue collected from C57BL/6 mice was put in the medium of ID8 cells, and co-cultured for 48 hours.

### ELISA

Cultured medium from ID8 and SK-OV-3 cells were collected. Total protein of cultured cells was extracted by using the cell lysis buffer (20 mM Tris-HCl (pH 7.5), 150 mM NaCl, 1 mM of EDTA, 1 mM EGTA, 1% Triton X-100, 2.5 mM sodium pyrophosphate, 1 mM beta-glycerophosphate, 1 mM Na_3_VO_4_, and protease inhibitor cocktail) and the concentration was measured by Pierce TM BCA Protein Assay Kit (Thermo-Fisher Scientific) for normalization. The ascites from ID8 orthotopic mouse OC model was collected at 60 days post-injection. Cultured medium and ascites were centrifuged at 1,000 ×g for 20 minutes at 4 ℃. The concentration of murine and human IL-18 in the supernatants were assayed by murine IL18 ELISA Kit (Wuxin Donglin Sci & Tech Development, Wuxi, Jiangsu, China) and human IL18 ELISA Kit (Wuxin Donglin Sci & Tech Development). The OD (450 nm) values were measured by Infinite M200 PRO (TECAN, Männedorf, Switzerland) microplate reader. The data were analyzed by Curve Expert (Hyams Development).

### Deep RNA sequencing

Total RNA of IL20RA-reconstituted and control ID8 cells were stimulated with IL-20 for 24 hours before harvested using TRIzol reagent for RNA extraction. The deep RNA sequencing was performed and analyzed on BGI seq500 platform (BGI-Shenzhen, China). KEGG pathway analyses were conducted and analyzed based on KEGG pathway database (http://www.genome.jp/kegg/).

### ChIP-qPCR

The ChIP was performed by using ChIP-IT® Express Kits according to the manufacturer’s instructions (Active Motif, San Diego, CA, USA). For immunoprecipitation, 7 μg of sheared chromatin DNA fragments were incubated with IgG or STAT3-specific antibodies. qPCR assay was used to detect the relative enrichment. The antibodies used in ChIP assay were listed on Supplemental Table 1. The primer sequences are listed in Supplemental Table 2.

### Study approval

Animal experiments were approved by Institutional Animal Care and Use Committee of Nankai University. The patients study received ethical approval from the ethics committee of Nankai University and Tianjin Center Hospital of Gynecology Obstetrics, and conformed to the principles embodied in the Declaration of Helsinki. All OC patients from Tianjin Center Hospital of Gynecology Obstetrics provided informed consent.

### Statistics

Prism 8.0 software (GraphPad Software, San Diego, CA, USA) was used for statistical analysis. Statistical parameters including the definitions and exact values of n, statistical test and statistical significance are reported in the Figures and Figure Legends. Quantitative data were presented as means ± SEM, and the differences between the groups were analyzed using the Student’s t-test. Survival curves were analyzed using the Kaplan–Meier method, and the log-rank test was used to calculate differences between the curves. Differences are considered statistically significant at *p < 0.05; **p < 0.01; ***p < 0.001; ns means no significance.

## Acknowledgements

This work was supported by grants from the National Natural Science Foundation of China (No. 81772974, No.81972882, No. 81874297) and Bilateral Inter-Governmental S&T Cooperation Project from Ministry of Science and Technology of China (2018YFE0114300).

## Author contributions

J.L., R.X. and Y.S. designed the study. J.L., X.Q0, X.W., J.H. and J.S. conducted the experiments and acquired data. J.L. and Y.Z. conducted the FACS gating and TAM analysis. Y.P. and X.C. conducted bioinformatics analysis. W.Z. and P.Q. acquired consent from patients, acquired samples, and collected clinical data. J.L., X.Q., J.S., T.L. and M.X. analyzed the data. J.L. and L.W. wrote the manuscript. R.X. and Y.S. supervised the research and edited the manuscript.

---

**Supplemental Figure 1.**
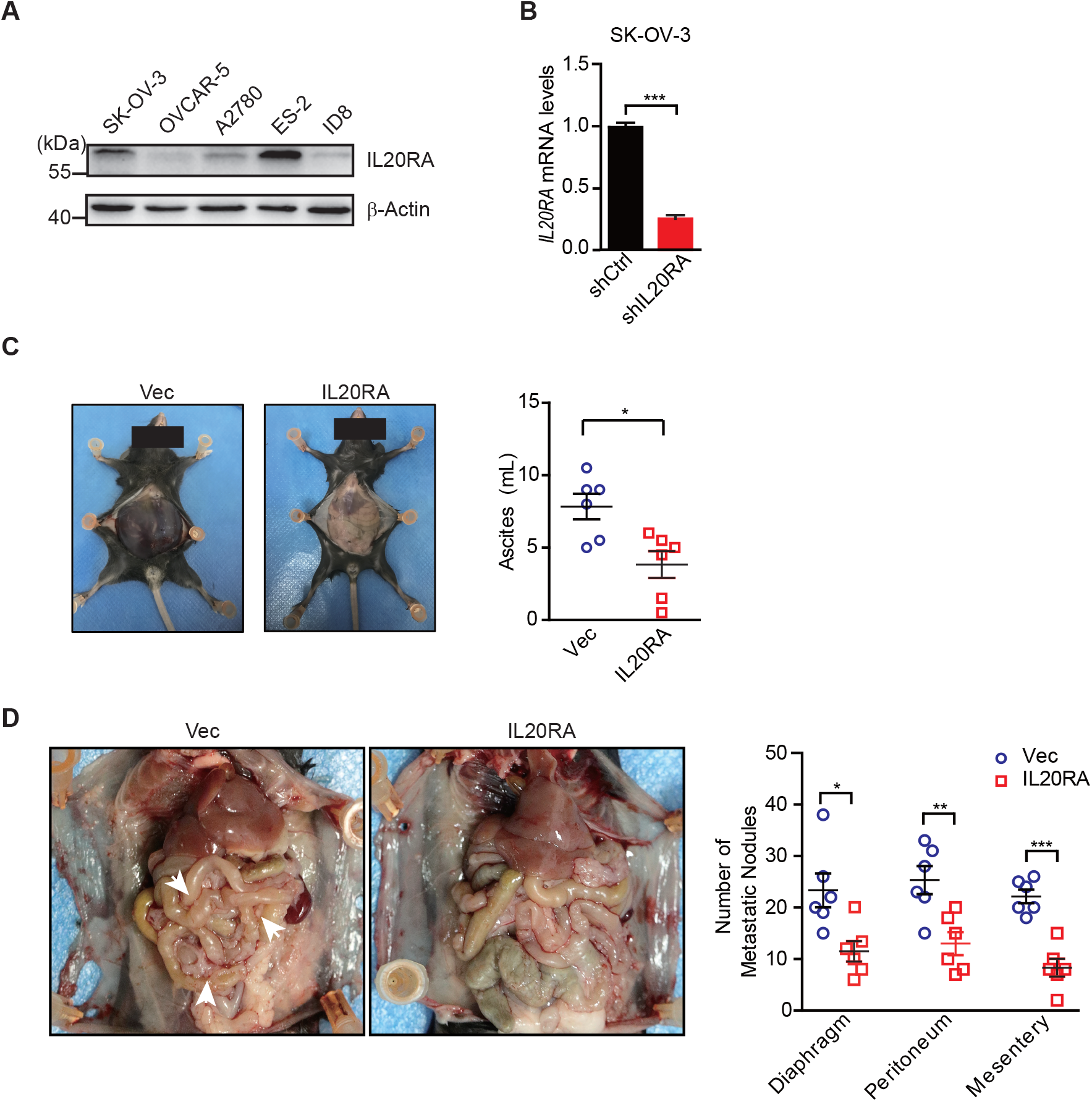
IL20RA suppresses the transcoelomic metastasis of OC in syngeneic intraperitoneal OC mouse model. **A.** Western blot analysis of IL20RA in different OC cells. **B.** qRT-PCR analysis of IL20RA in SK-OV-3 cells transfected with shCtrl or shIL20RA (means ± SEM, ***P<0.001, by unpaired two-sided student’s t-test). **C-D.** Representative images of C57BL/6 mice at 45 days after direct intraperitoneal injection of IL20RA-reconstituted or control (Vec) ID8 cells (C and D, left panels). The ascites volumes and numbers of metastatic nodules on the surfaces of diaphragm, peritoneum and mesentery were quantified (n = 6 mice for each group). Data are shown as means ± SEM, *P < 0.05, **P < 0.01, ***P < 0.001, by unpaired two-sided student’s t-test.

**Supplemental Figure 2.**
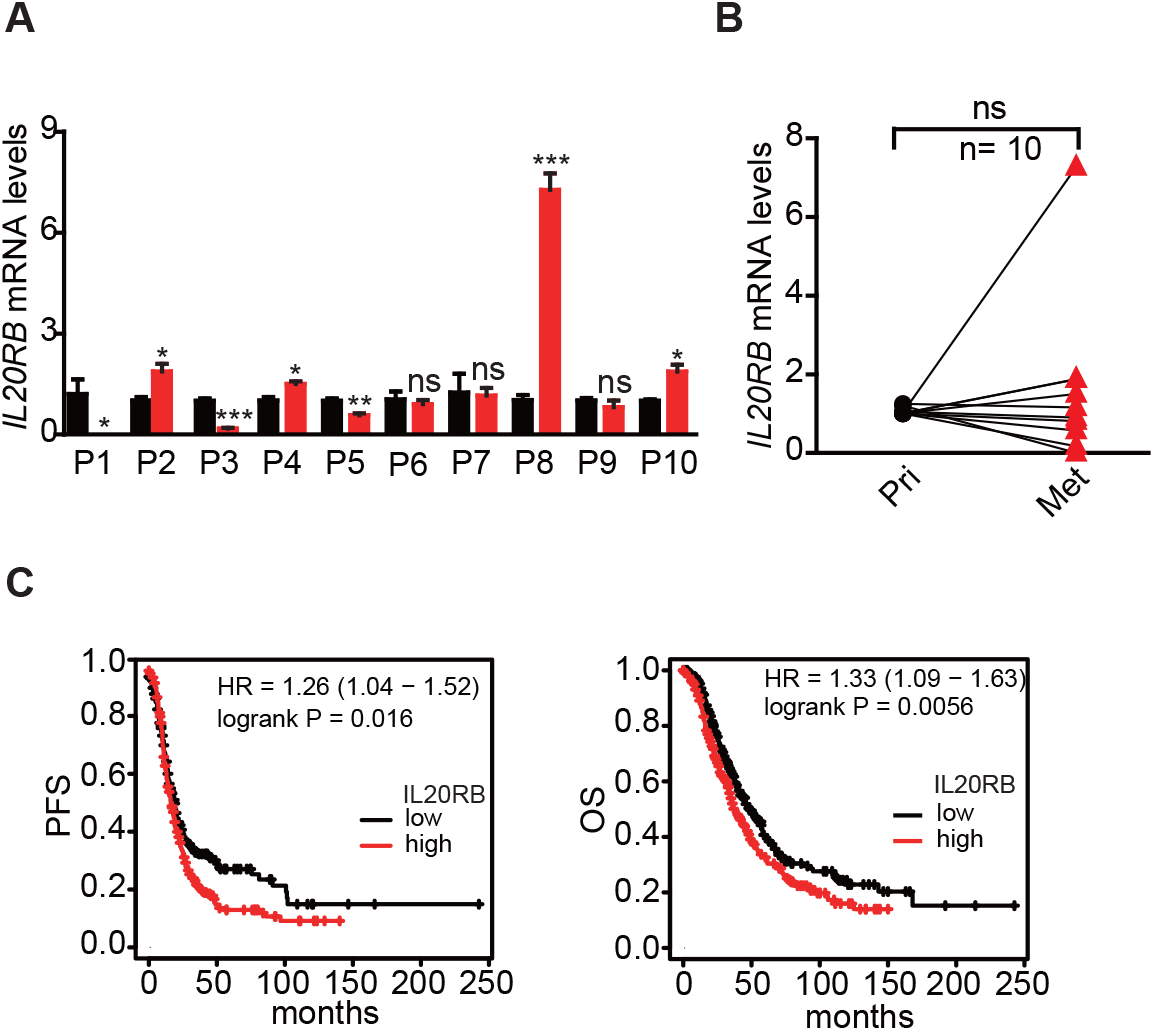
IL20RB expression in human OC tissues and its correlation with clinical outcome of OC patients. **A-B.** qRT-PCR analysis of IL20RB in human primary OC tissue and paired peritoneal metastases (**A**, data are plotted as means ± SEM from three independent measurements, *P < 0.05, **P < 0.01, ***P < 0.001, ns not significant, by unpaired two-sided student’s t-test). The comparison of the IL20RB levels in these two groups are statistically analyzed in (**B**, ns not significant, by unpaired two-sided student’s t-test). **C.** Kaplan-Meier survival plot to show the progression free survival (PFS) and overall survival (OS) of OC patients with different IL20RB expression.

**Supplemental Figure 3.**
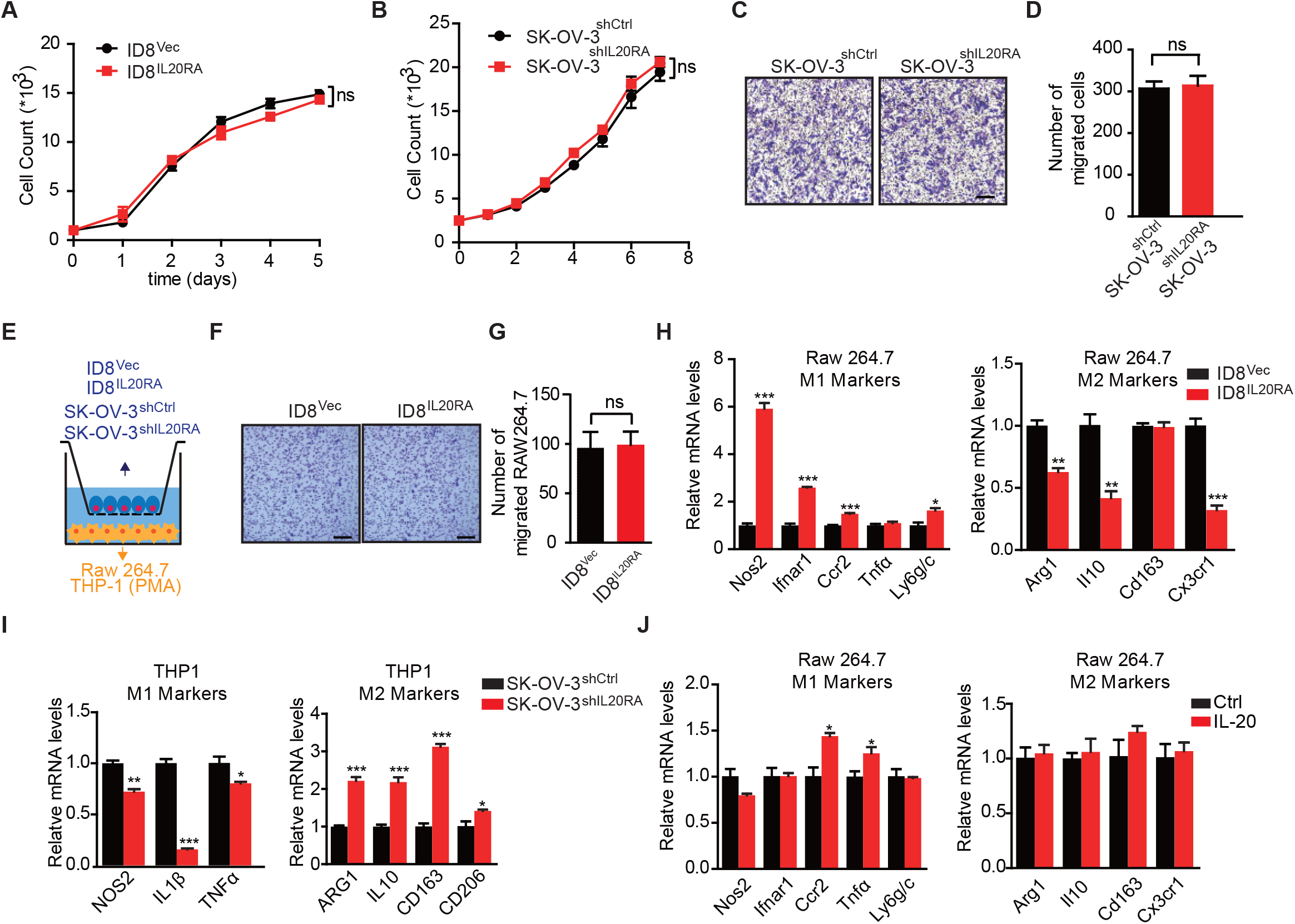
*In vitro* co-culture experiments to show the IL20RA-mediated crosstalk between OC cells and macrophages for their polarization. **A.** Growth curves of IL20RA-reconstituted or control (Vec) ID8 cells under the stimulation of IL-20. **B**. Growth curves of shIL20RA- or shCtrl-transfected SK-OV-3 cells under the stimulation of IL-20. **C-D.** Transwell cell migration assay of shIL20RA- or shCtrl-transfected SK-OV-3 cells in the presence of IL-20, which were quantified in (**D**, shown as means ± SEM from three independent experiments, ns not significant, by unpaired two-sided student’s t-test). Scale bar: 50 μm. **E.** Schematic of the *in vitro* co-culture experiments. **F-G.** Transwell migration assay to show the migrated RAW 264.7 cells co-cultured with IL20RA-reconstituted or control (Vec) ID8 cells (**F**), which were quantified in (**G**, shown as means ± SEM from three independent experiments, ns not significant, by unpaired two-sided student’s t-test). Scale bar: 50 μm. **H.** qRT-PCR analysis of macrophage marker genes in BMDM co-cultured with IL20RA-reconstituted or control (Vec) ID8 cells for 72 h. **I.** qRT-PCR analysis of macrophage marker genes in THP-1-derived macrophages co-cultured with shIL20RA-or shCtrl-transfected SK-OV-3 cells for 72 h. **J.** qRT-PCR analysis of macrophage marker genes in RAW 264.7 cells stimulated with IL-20 protein for 72 h. All the qRT-PCR data are shown as means ± SEM from three independent experiments, *P < 0.05, **P < 0.01, ***P < 0.001, by unpaired two-sided student’s t-test.

**Supplemental Figure 4.**
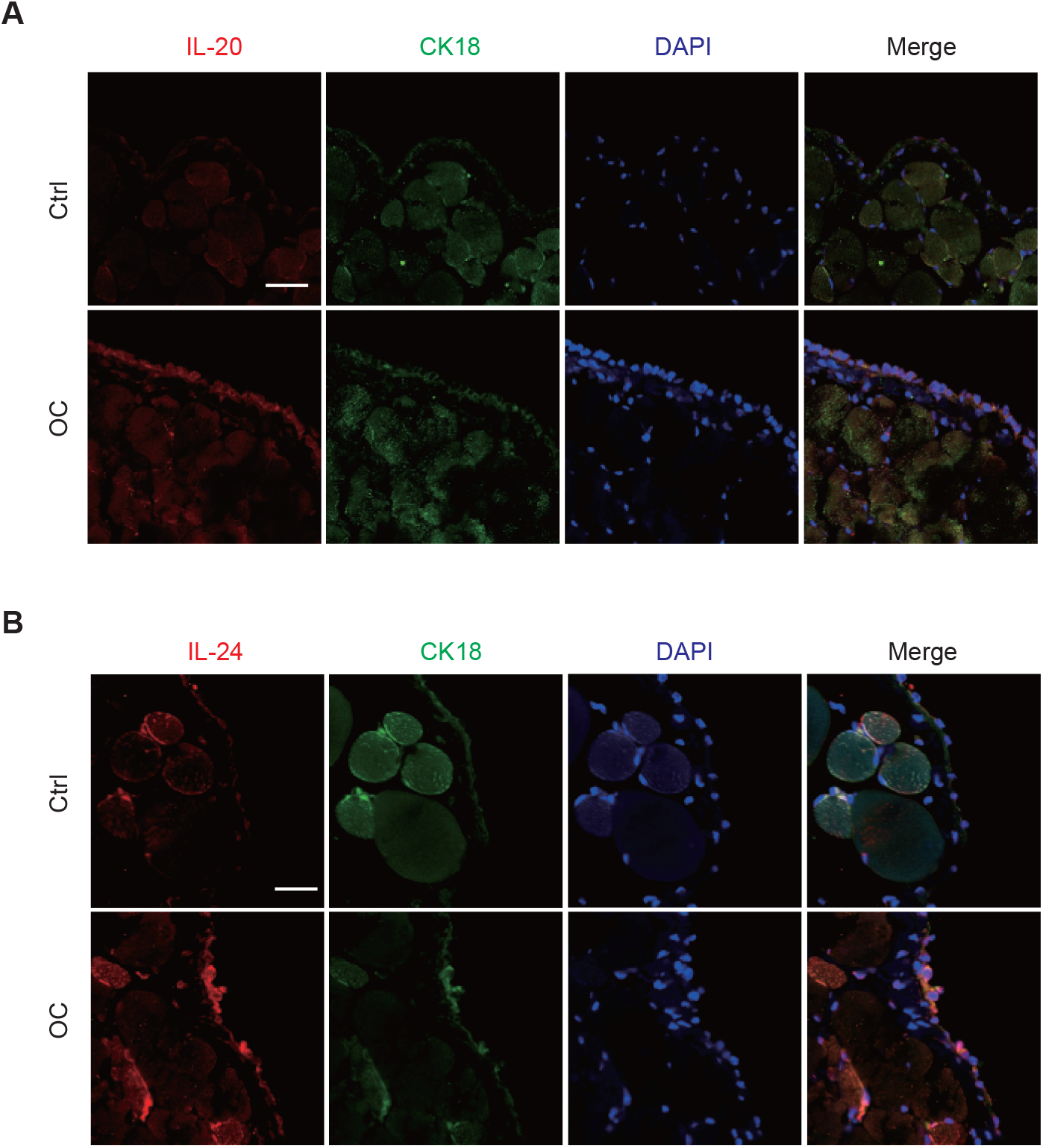
Mesothelial cells in peritoneum are the major source of IL-20 and IL-24 when challenged by OC. **A-B.** IF staining of IL-20/IL-24 (red), cytokeratin Ck18 (green) and DAPI (blue) in peritoneum dissected from mice with intraperitoneal injection of ID8 cells (OC) or PBS control (Ctrl) 9 days before. Scale bar: 30 μm.

**Supplemental Figure 5.**
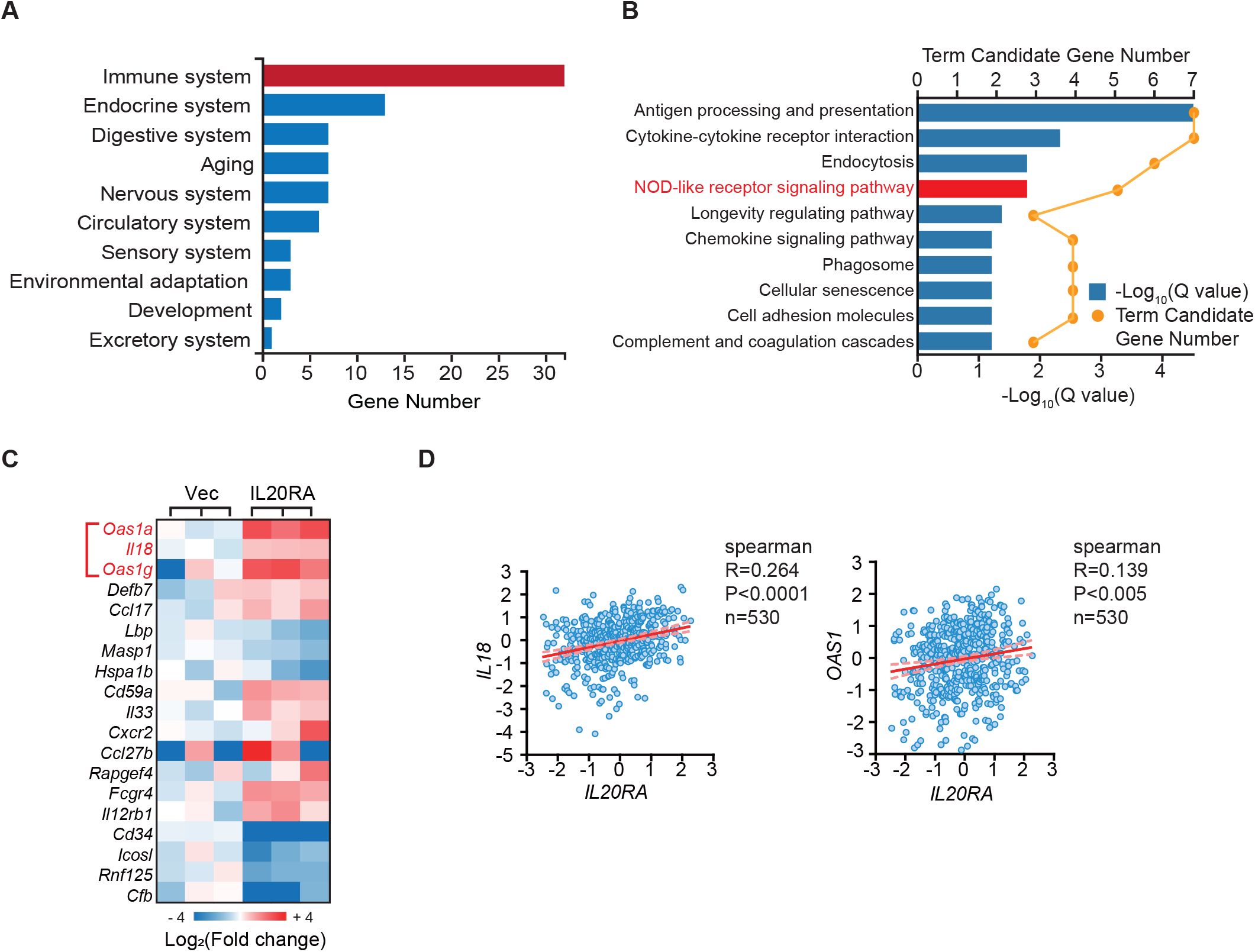
KEGG analysis showes IL-20 activates the OAS1/RNase L-NLR/IL-18 signaling. **A.** KEGG gene set (based on organism systems) enrichment analysis on significantly changed genes in IL20RA-reconstituted ID8 cells. **B**. KEGG pathway enrichment analysis on significantly changed, immune-related genes in IL20RA-reconstituted ID8 cells. **C.** Heatmap of differentially expressed, immune-related genes in IL20RA-reconstituted ID8 cells. **D.** The Spearman’s correlation analysis on the expression of *IL20RA*, *OAS1* and *IL18* in OC patients..

**Supplemental Figure 6.**
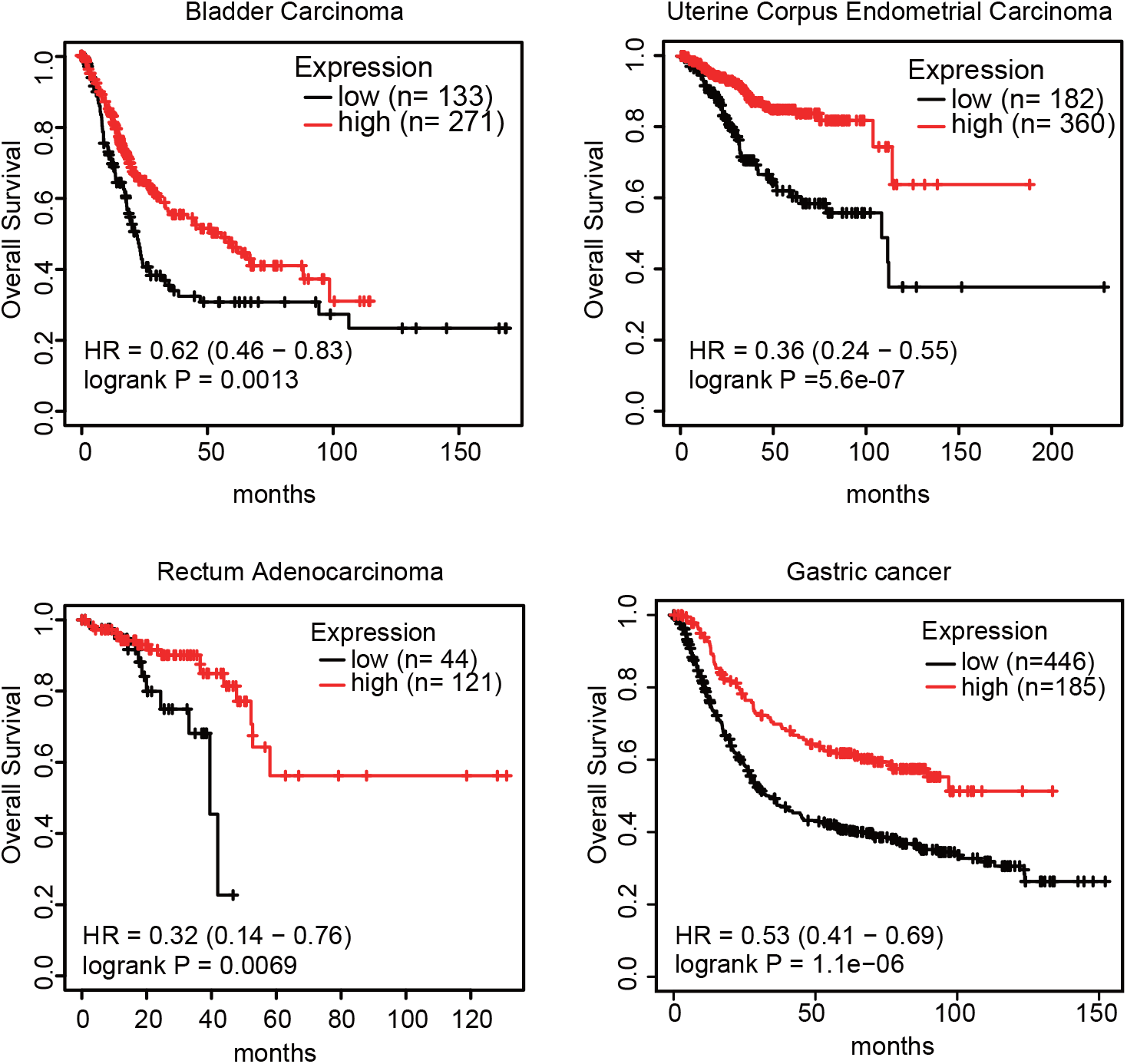
IL20RA expression is positively correlated with the clinical outcome of patients in many types of cancers. Kaplan-Meier survival plots to show the correlations of different IL20RA expression with the overall survival in patients of bladder carcinoma, uterine corpus endometrial carcinoma, rectum adenocarcinoma and gastric cancer.

**Supplemental Table 1.**
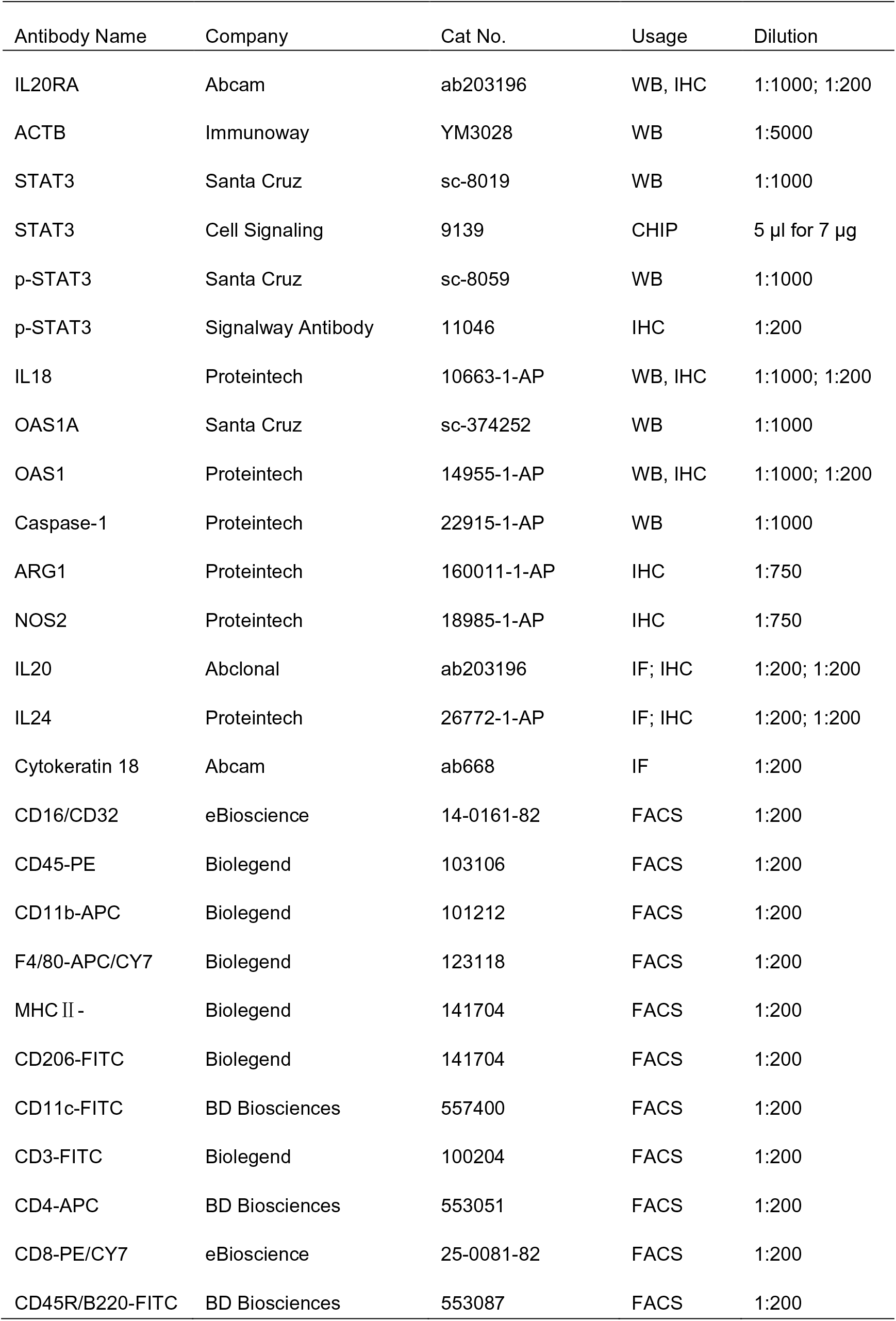
List of antibodies

| Antibody Name | Company | Cat No. | Usage | Dilution |
| --- | --- | --- | --- | --- |
| IL20RA | Abcam | ab203196 | WB, IHC | 1:1000; 1:200 |
| ACTB | Immunoway | YM3028 | WB | 1:5000 |
| STAT3 | Santa Cruz | sc-8019 | WB | 1:1000 |
| STAT3 | Cell Signaling | 9139 | CHIP | 5 µl for 7 µg |
| p-STAT3 | Santa Cruz | sc-8059 | WB | 1:1000 |
| p-STAT3 | Signalway Antibody | 11046 | IHC | 1:200 |
| IL18 | Proteintech | 10663-1-AP | WB, IHC | 1:1000; 1:200 |
| OAS1A | Santa Cruz | sc-374252 | WB | 1:1000 |
| OAS1 | Proteintech | 14955-1-AP | WB, IHC | 1:1000; 1:200 |
| Caspase-1 | Proteintech | 22915-1-AP | WB | 1:1000 |
| ARG1 | Proteintech | 160011-1-AP | IHC | 1:750 |
| NOS2 | Proteintech | 18985-1-AP | IHC | 1:750 |
| IL20 | Abclonal | ab203196 | IF; IHC | 1:200; 1:200 |
| IL24 | Proteintech | 26772-1-AP | IF; IHC | 1:200; 1:200 |
| Cytokeratin 18 | Abcam | ab668 | IF | 1:200 |
| CD16/CD32 | eBioscience | 14-0161-82 | FACS | 1:200 |
| CD45-PE | Biolegend | 103106 | FACS | 1:200 |
| CD11b-APC | Biolegend | 101212 | FACS | 1:200 |
| F4/80-APC/CY7 | Biolegend | 123118 | FACS | 1:200 |
| MHC II - | Biolegend | 141704 | FACS | 1:200 |
| CD206-FITC | Biolegend | 141704 | FACS | 1:200 |
| CD11c-FITC | BD Biosciences | 557400 | FACS | 1:200 |
| CD3-FITC | Biolegend | 100204 | FACS | 1:200 |
| CD4-APC | BD Biosciences | 553051 | FACS | 1:200 |
| CD8-PE/CY7 | eBioscience | 25-0081-82 | FACS | 1:200 |
| CD45R/B220-FITC | BD Biosciences | 553087 | FACS | 1:200 |

**Supplemental Table 2.**
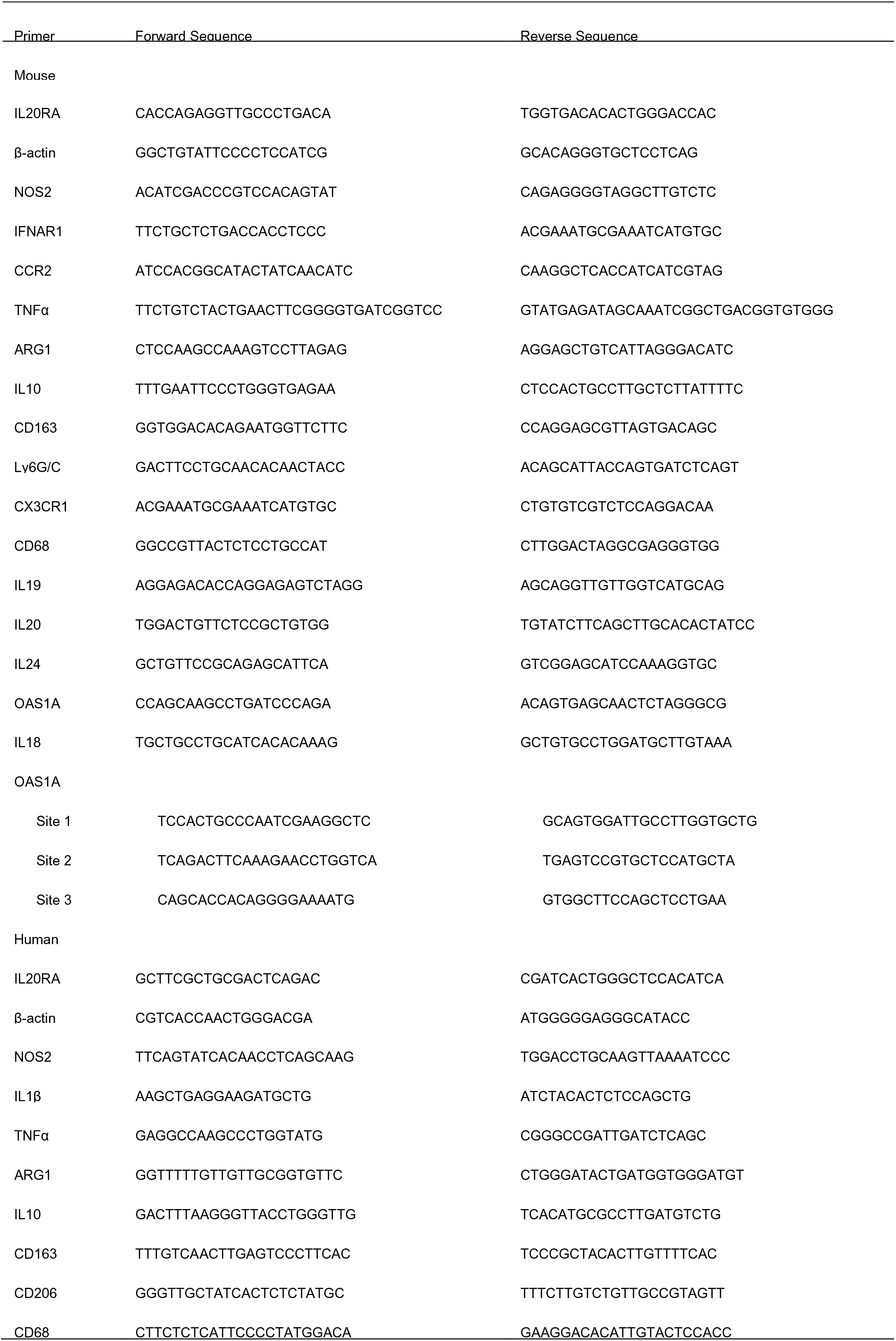
Primes for qPCR (human and mouse)

## References

Ahmed, N., & Stenvers, K. L. (2013). Getting to know ovarian cancer ascites: opportunities for targeted therapy-based translational research. Frontiers in Oncology, 3, 256. https://doi.org/10.3389/fonc.2013.00256

Aziz, M., Agarwal, K., Dasari, S., & Mitra, A. A. K. (2019). Productive Cross-Talk with the Microenvironment: A Critical Step in Ovarian Cancer Metastasis. Cancers, 11(10). https://doi.org/10.3390/cancers11101608

Banerjee, S. (2016). RNase L and the NLRP3-inflammasome: An old merchant in a new trade. Cytokine and Growth Factor Reviews, 29, 63–70. https://doi.org/10.1016/j.cytogfr.2016.02.008

Chakrabarti, A., Banerjee, S., Franchi, L., Loo, Y. M., Gale, M., Jr., Nunez, G., & Silverman, R. H. (2015). RNase L activates the NLRP3 inflammasome during viral infections. Cell Host Microbe, 17(4), 466–477. https://doi.org/10.1016/j.chom.2015.02.010

Chang, A., Chen, Y., Shen, W., Gao, R., Zhou, W., Yang, S., Liu, Y., Luo, Y., Chuang, T. H., Sun, P., Liu, C., & Xiang, R. (2015). Ifit1 Protects Against Lipopolysaccharide and D-galactosamine-Induced Fatal Hepatitis by Inhibiting Activation of the JNK Pathway. Journal of Infectious Diseases, 212(9), 1509–1520. https://doi.org/10.1093/infdis/jiv221

Colmont, C. S., Raby, A. C., Dioszeghy, V., Lebouder, E., Foster, T. L., Jones, S. A., Labeta, M. O., Fielding, C. A., & Topley, N. (2011). Human peritoneal mesothelial cells respond to bacterial ligands through a specific subset of Toll-like receptors. Nephrology, Dialysis, Transplantation, 26(12), 4079–4090. https://doi.org/10.1093/ndt/gfr217

Coughlin, C. M., Salhany, K. E., Wysocka, M., Aruga, E., Kurzawa, H., Chang, A. E., Hunter, C. A., Fox, J. C., Trinchieri, G., & Lee, W. M. (1998). Interleukin-12 and interleukin-18 synergistically induce murine tumor regression which involves inhibition of angiogenesis. Journal of Clinical Investigation, 101(6), 1441–1452. https://doi.org/10.1172/JCI1555

Diaz Osterman, C. J., Ozmadenci, D., Kleinschmidt, E. G., Taylor, K. N., Barrie, A. M., Jiang, S., Bean, L. M., Sulzmaier, F. J., Jean, C., Tancioni, I., Anderson, K., Uryu, S., Cordasco, E. A., Li, J., Chen, X. L., Fu, G., Ojalill, M., Rappu, P., Heino, J., Mark, A. M., Xu, G., Fisch, K. M., Kolev, V. N., Weaver, D. T., Pachter, J. A., Gyorffy, B., McHale, M. T., Connolly, D. C., Molinolo, A., Stupack, D. G., & Schlaepfer, D. D. (2019). FAK activity sustains intrinsic and acquired ovarian cancer resistance to platinum chemotherapy. Elife, 8. https://doi.org/10.7554/eLife.47327

Fabbi, M., Carbotti, G., & Ferrini, S. (2015). Context-dependent role of IL-18 in cancer biology and counter-regulation by IL-18BP. Journal of Leukocyte Biology, 97(4), 665–675. https://doi.org/10.1189/jlb.5RU0714-360RR

Farley, J., Ozbun, L. L., & Birrer, M. J. (2008). Genomic analysis of epithelial ovarian cancer. Cell Research, 18(5), 538–548. https://doi.org/10.1038/cr.2008.52

Feng, Q., Li, P., Salamanca, C., Huntsman, D., Leung, P. C., & Auersperg, N. (2005). Caspase-1alpha is down-regulated in human ovarian cancer cells and the overexpression of caspase-1alpha induces apoptosis. Cancer Research, 65(19), 8591–8596. https://doi.org/10.1158/0008-5472.CAN-05-0239

Fisher, P. B., Sarkar, D., Lebedeva, I. V., Emdad, L., Gupta, P., Sauane, M., Su, Z. Z., Grant, S., Dent, P., Curiel, D. T., Senzer, N., & Nemunaitis, J. (2007). Melanoma differentiation associated gene-7/interleukin-24 (mda-7/IL-24): novel gene therapeutic for metastatic melanoma. Toxicology and Applied Pharmacology, 224(3), 300–307. https://doi.org/10.1016/j.taap.2006.11.021

Gopalan, B., Shanker, M., Chada, S., & Ramesh, R. (2007). MDA-7/IL-24 suppresses human ovarian carcinoma growth in vitro and in vivo. Molecular Cancer, 6, 11. https://doi.org/10.1186/1476-4598-6-11

Gyorffy, B., Lanczky, A., & Szallasi, Z. (2012). Implementing an online tool for genome-wide validation of survival-associated biomarkers in ovarian-cancer using microarray data from 1287 patients. Endocrine-Related Cancer, 19(2), 197–208. https://doi.org/10.1530/ERC-11-0329

Hsu, Y. H., Hsing, C. H., Li, C. F., Chan, C. H., Chang, M. C., Yan, J. J., & Chang, M. S. (2012). Anti-IL-20 monoclonal antibody suppresses breast cancer progression and bone osteolysis in murine models. Journal of Immunology, 188(4), 1981–1991. https://doi.org/10.4049/jimmunol.1102843

Huang, K., Liu, X., Li, Y., Wang, Q., Zhou, J., Wang, Y., Dong, F., Yang, C., Sun, Z., Fang, C., Liu, C., Tan, Y., Wu, X., Jiang, T., & Kang, C. (2019). Genome-Wide CRISPR-Cas9 Screening Identifies NF-kappaB/E2F6 Responsible for EGFRvIII-Associated Temozolomide Resistance in Glioblastoma. Adv Sci (Weinh), 6(17), 1900782. https://doi.org/10.1002/advs.201900782

Isaza-Restrepo, A., Martin-Saavedra, J. S., Velez-Leal, J. L., Vargas-Barato, F., & Riveros-Duenas, R. (2018). The Peritoneum: Beyond the Tissue - A Review. Frontiers in Physiology, 9, 738. https://doi.org/10.3389/fphys.2018.00738

Iwanicki, M. P., Davidowitz, R. A., Ng, M. R., Besser, A., Muranen, T., Merritt, M., Danuser, G., Ince, T. A., & Brugge, J. S. (2011). Ovarian cancer spheroids use myosin-generated force to clear the mesothelium. Cancer Discovery, 1(2), 144–157. https://doi.org/10.1158/2159-8274.CD-11-0010

Joung, J., Konermann, S., Gootenberg, J. S., Abudayyeh, O. O., Platt, R. J., Brigham, M. D., Sanjana, N. E., & Zhang, F. (2017). Genome-scale CRISPR-Cas9 knockout and transcriptional activation screening. Nature Protocols, 12(4), 828–863. https://doi.org/10.1038/nprot.2017.016

Kato, S., Yuzawa, Y., Tsuboi, N., Maruyama, S., Morita, Y., Matsuguchi, T., & Matsuo, S. (2004). Endotoxin-induced chemokine expression in murine peritoneal mesothelial cells: the role of toll-like receptor 4. Journal of the American Society of Nephrology, 15(5), 1289–1299. http://www.ncbi.nlm.nih.gov/pubmed/15100369

Kenny, H. A., Krausz, T., Yamada, S. D., & Lengyel, E. (2007). Use of a novel 3D culture model to elucidate the role of mesothelial cells, fibroblasts and extra-cellular matrices on adhesion and invasion of ovarian cancer cells to the omentum. International Journal of Cancer, 121(7), 1463–1472. https://doi.org/10.1002/ijc.22874

Kenny, H. A., Nieman, K. M., Mitra, A. K., & Lengyel, E. (2011). The first line of intra-abdominal metastatic attack: breaching the mesothelial cell layer. Cancer Discovery, 1(2), 100–102. https://doi.org/10.1158/2159-8290.CD-11-0117

Kipps, E., Tan, D. S., & Kaye, S. B. (2013). Meeting the challenge of ascites in ovarian cancer: new avenues for therapy and research. Nature Reviews: Cancer, 13(4), 273–282. https://doi.org/10.1038/nrc3432

Lee, S. J., Cho, S. C., Lee, E. J., Kim, S., Lee, S. B., Lim, J. H., Choi, Y. H., Kim, W. J., & Moon, S. K. (2013). Interleukin-20 promotes migration of bladder cancer cells through extracellular signal-regulated kinase (ERK)-mediated MMP-9 protein expression leading to nuclear factor (NF-kappaB) activation by inducing the up-regulation of p21(WAF1) protein expression. Journal of Biological Chemistry, 288(8), 5539–5552. https://doi.org/10.1074/jbc.M112.410233

Mikula-Pietrasik, J., Uruski, P., Tykarski, A., & Ksiazek, K. (2018). The peritoneal “soil” for a cancerous “seed”: a comprehensive review of the pathogenesis of intraperitoneal cancer metastases. Cellular and Molecular Life Sciences, 75(3), 509–525. https://doi.org/10.1007/s00018-017-2663-1

Motohara, T., Masuda, K., Morotti, M., Zheng, Y., El-Sahhar, S., Chong, K. Y., Wietek, N., Alsaadi, A., Karaminejadranjbar, M., Hu, Z., Artibani, M., Gonzalez, L. S., Katabuchi, H., Saya, H., & Ahmed, A. A. (2019). An evolving story of the metastatic voyage of ovarian cancer cells: cellular and molecular orchestration of the adipose-rich metastatic microenvironment. Oncogene, 38(16), 2885–2898. https://doi.org/10.1038/s41388-018-0637-x

Musteanu, M., Blaas, L., Mair, M., Schlederer, M., Bilban, M., Tauber, S., Esterbauer, H., Mueller, M., Casanova, E., Kenner, L., Poli, V., & Eferl, R. (2010). Stat3 is a negative regulator of intestinal tumor progression in Apc(Min) mice. Gastroenterology, 138(3), 1003–1011 e1001-1005. https://doi.org/10.1053/j.gastro.2009.11.049

Ng, S. R., Rideout, W. M., 3rd, Akama-Garren, E. H., Bhutkar, A., Mercer, K. L., Schenkel, J. M., Bronson, R. T., & Jacks, T. (2020). CRISPR-mediated modeling and functional validation of candidate tumor suppressor genes in small cell lung cancer. Proceedings of the National Academy of Sciences of the United States of America, 117(1), 513–521. https://doi.org/10.1073/pnas.1821893117

Niedbala, M. J., Crickard, K., & Bernacki, R. J. (1985). Interactions of human ovarian tumor cells with human mesothelial cells grown on extracellular matrix. An in vitro model system for studying tumor cell adhesion and invasion. Experimental Cell Research, 160(2), 499–513. https://doi.org/10.1016/0014-4827(85)90197-1

Nishio, S., Yamada, N., Ohyama, H., Yamanegi, K., Nakasho, K., Hata, M., Nakamura, Y., Fukunaga, S., Futani, H., Yoshiya, S., Ueda, H., Taniguchi, M., Okamura, H., & Terada, N. (2008). Enhanced suppression of pulmonary metastasis of malignant melanoma cells by combined administration of alpha-galactosylceramide and interleukin-18. Cancer Science, 99(1), 113–120. https://doi.org/10.1111/j.1349-7006.2007.00636.x

Orengo, A. M., Fabbi, M., Miglietta, L., Andreani, C., Bruzzone, M., Puppo, A., Cristoforoni, P., Centurioni, M. G., Gualco, M., Salvi, S., Boccardo, S., Truini, M., Piazza, T., Canevari, S., Mezzanzanica, D., & Ferrini, S. (2011). Interleukin (IL)-18, a biomarker of human ovarian carcinoma, is predominantly released as biologically inactive precursor. International Journal of Cancer, 129(5), 1116–1125. https://doi.org/10.1002/ijc.25757

Parayath, N. N., Gandham, S. K., Leslie, F., & Amiji, M. M. (2019). Improved anti-tumor efficacy of paclitaxel in combination with MicroRNA-125b-based tumor-associated macrophage repolarization in epithelial ovarian cancer. Cancer Letters, 461, 1–9. https://doi.org/10.1016/j.canlet.2019.07.002

Park, J. H., Kim, Y. G., Shaw, M., Kanneganti, T. D., Fujimoto, Y., Fukase, K., Inohara, N., & Nunez, G. (2007). Nod1/RICK and TLR signaling regulate chemokine and antimicrobial innate immune responses in mesothelial cells. Journal of Immunology, 179(1), 514–521. https://doi.org/10.4049/jimmunol.179.1.514

Pestka, S., Krause, C. D., Sarkar, D., Walter, M. R., Shi, Y., & Fisher, P. B. (2004). Interleukin-10 and related cytokines and receptors. Annual Review of Immunology, 22, 929–979. https://doi.org/10.1146/annurev.immunol.22.012703.104622

Pradhan, A. K., Bhoopathi, P., Talukdar, S., Shen, X. N., Emdad, L., Das, S. K., Sarkar, D., & Fisher, P. B. (2018). Recombinant MDA-7/IL24 Suppresses Prostate Cancer Bone Metastasis through Downregulation of the Akt/Mcl-1 Pathway. Molecular Cancer Therapeutics, 17(9), 1951–1960. https://doi.org/10.1158/1535-7163.MCT-17-1002

Reinartz, S., Schumann, T., Finkernagel, F., Wortmann, A., Jansen, J. M., Meissner, W., Krause, M., Schworer, A. M., Wagner, U., Muller-Brusselbach, S., & Muller, R. (2014). Mixed-polarization phenotype of ascites-associated macrophages in human ovarian carcinoma: correlation of CD163 expression, cytokine levels and early relapse. International Journal of Cancer, 134(1), 32–42. https://doi.org/10.1002/ijc.28335

Robertson, M. J., Kline, J., Struemper, H., Koch, K. M., Bauman, J. W., Gardner, O. S., Murray, S. C., Germaschewski, F., Weisenbach, J., Jonak, Z., & Toso, J. F. (2013). A dose-escalation study of recombinant human interleukin-18 in combination with rituximab in patients with non-Hodgkin lymphoma. Journal of Immunotherapy, 36(6), 331–341. https://doi.org/10.1097/CJI.0b013e31829d7e2e

Robertson, M. J., Mier, J. W., Logan, T., Atkins, M., Koon, H., Koch, K. M., Kathman, S., Pandite, L. N., Oei, C., Kirby, L. C., Jewell, R. C., Bell, W. N., Thurmond, L. M., Weisenbach, J., Roberts, S., & Dar, M. M. (2006). Clinical and biological effects of recombinant human interleukin-18 administered by intravenous infusion to patients with advanced cancer. Clinical Cancer Research, 12(14 Pt 1), 4265–4273. https://doi.org/10.1158/1078-0432.CCR-06-0121

Rolny, C., Mazzone, M., Tugues, S., Laoui, D., Johansson, I., Coulon, C., Squadrito, M. L., Segura, I., Li, X., Knevels, E., Costa, S., Vinckier, S., Dresselaer, T., Akerud, P., De Mol, M., Salomaki, H., Phillipson, M., Wyns, S., Larsson, E., Buysschaert, I., Botling, J., Himmelreich, U., Van Ginderachter, J. A., De Palma, M., Dewerchin, M., Claesson-Welsh, L., & Carmeliet, P. (2011). HRG inhibits tumor growth and metastasis by inducing macrophage polarization and vessel normalization through downregulation of PlGF. Cancer Cell, 19(1), 31–44. https://doi.org/10.1016/j.ccr.2010.11.009

Rutz, S., Wang, X., & Ouyang, W. (2014). The IL-20 subfamily of cytokines--from host defence to tissue homeostasis. Nature Reviews: Immunology, 14(12), 783–795. https://doi.org/10.1038/nri3766

Salcedo, R., Worschech, A., Cardone, M., Jones, Y., Gyulai, Z., Dai, R. M., Wang, E., Ma, W., Haines, D., O’HUigin, C., Marincola, F. M., & Trinchieri, G. (2010). MyD88-mediated signaling prevents development of adenocarcinomas of the colon: role of interleukin 18. Journal of Experimental Medicine, 207(8), 1625–1636. https://doi.org/10.1084/jem.20100199

Sandoval, P., Jimenez-Heffernan, J. A., Rynne-Vidal, A., Perez-Lozano, M. L., Gilsanz, A., Ruiz-Carpio, V., Reyes, R., Garcia-Bordas, J., Stamatakis, K., Dotor, J., Majano, P. L., Fresno, M., Cabanas, C., & Lopez-Cabrera, M. (2013). Carcinoma-associated fibroblasts derive from mesothelial cells via mesothelial-to-mesenchymal transition in peritoneal metastasis. Journal of Pathology, 231(4), 517–531. https://doi.org/10.1002/path.4281

Shalem, O., Sanjana, N. E., Hartenian, E., Shi, X., Scott, D. A., Mikkelson, T., Heckl, D., Ebert, B. L., Root, D. E., Doench, J. G., & Zhang, F. (2014). Genome-scale CRISPR-Cas9 knockout screening in human cells. Science, 343(6166), 84–87. https://doi.org/10.1126/science.1247005

Siegel, R. L., Miller, K. D., & Jemal, A. (2020). Cancer statistics, 2020. CA: A Cancer Journal for Clinicians, 70(1), 7–30. https://doi.org/10.3322/caac.21590

Tan, D. S., Agarwal, R., & Kaye, S. B. (2006). Mechanisms of transcoelomic metastasis in ovarian cancer. Lancet Oncology, 7(11), 925–934. https://doi.org/10.1016/S1470-2045(06)70939-1

Tarhini, A. A., Millward, M., Mainwaring, P., Kefford, R., Logan, T., Pavlick, A., Kathman, S. J., Laubscher, K. H., Dar, M. M., & Kirkwood, J. M. (2009). A phase 2, randomized study of SB-485232, rhIL-18, in patients with previously untreated metastatic melanoma. Cancer, 115(4), 859–868. https://doi.org/10.1002/cncr.24100

Thibault, B., Castells, M., Delord, J. P., & Couderc, B. (2014). Ovarian cancer microenvironment: implications for cancer dissemination and chemoresistance acquisition. Cancer and Metastasis Reviews, 33(1), 17–39. https://doi.org/10.1007/s10555-013-9456-2

Torre, L. A., Trabert, B., DeSantis, C. E., Miller, K. D., Samimi, G., Runowicz, C. D., Gaudet, M. M., Jemal, A., & Siegel, R. L. (2018). Ovarian cancer statistics, 2018. CA: A Cancer Journal for Clinicians, 68(4), 284–296. https://doi.org/10.3322/caac.21456

Vaughan, S., Coward, J. I., Bast, R. C., Jr., Berchuck, A., Berek, J. S., Brenton, J. D., Coukos, G., Crum, C. C., Drapkin, R., Etemadmoghadam, D., Friedlander, M., Gabra, H., Kaye, S. B., Lord, C. J., Lengyel, E., Levine, D. A., McNeish, I. A., Menon, U., Mills, G. B., Nephew, K. P., Oza, A. M., Sood, A. K., Stronach, E. A., Walczak, H., Bowtell, D. D., & Balkwill, F. R. (2011). Rethinking ovarian cancer: recommendations for improving outcomes. Nature Reviews: Cancer, 11(10), 719–725. https://doi.org/10.1038/nrc3144

Wang, Z. Y., Gaggero, A., Rubartelli, A., Rosso, O., Miotti, S., Mezzanzanica, D., Canevari, S., & Ferrini, S. (2002). Expression of interleukin-18 in human ovarian carcinoma and normal ovarian epithelium: evidence for defective processing in tumor cells. International Journal of Cancer, 98(6), 873–878. https://doi.org/10.1002/ijc.10268

Ward, K. K., Tancioni, I., Lawson, C., Miller, N. L., Jean, C., Chen, X. L., Uryu, S., Kim, J., Tarin, D., Stupack, D. G., Plaxe, S. C., & Schlaepfer, D. D. (2013). Inhibition of focal adhesion kinase (FAK) activity prevents anchorage-independent ovarian carcinoma cell growth and tumor progression. Clinical and Experimental Metastasis, 30(5), 579–594. https://doi.org/10.1007/s10585-012-9562-5

Whitaker, E. L., Filippov, V. A., & Duerksen-Hughes, P. J. (2012). Interleukin 24: mechanisms and therapeutic potential of an anti-cancer gene. Cytokine and Growth Factor Reviews, 23(6), 323–331. https://doi.org/10.1016/j.cytogfr.2012.08.004

Worzfeld, T., Pogge von Strandmann, E., Huber, M., Adhikary, T., Wagner, U., Reinartz, S., & Muller, R. (2017). The Unique Molecular and Cellular Microenvironment of Ovarian Cancer. Frontiers in Oncology, 7, 24. https://doi.org/10.3389/fonc.2017.00024

Yao, V., Platell, C., & Hall, J. C. (2004). Peritoneal mesothelial cells produce inflammatory related cytokines. ANZ Journal of Surgery, 74(11), 997–1002. https://doi.org/10.1111/j.1445-1433.2004.03220.x

You, W., Tang, Q., Zhang, C., Wu, J., Gu, C., Wu, Z., & Li, X. (2013). IL-26 promotes the proliferation and survival of human gastric cancer cells by regulating the balance of STAT1 and STAT3 activation. PloS One, 8(5), e63588. https://doi.org/10.1371/journal.pone.0063588

Zaki, M. H., Vogel, P., Body-Malapel, M., Lamkanfi, M., & Kanneganti, T. D. (2010). IL-18 production downstream of the Nlrp3 inflammasome confers protection against colorectal tumor formation. Journal of Immunology, 185(8), 4912–4920. https://doi.org/10.4049/jimmunol.1002046

Zhang, M., He, Y., Sun, X., Li, Q., Wang, W., Zhao, A., & Di, W. (2014). A high M1/M2 ratio of tumor-associated macrophages is associated with extended survival in ovarian cancer patients. J Ovarian Res, 7, 19. https://doi.org/10.1186/1757-2215-7-19

